# Noise robustness and metabolic load determine the principles of central dogma regulation

**DOI:** 10.1101/2023.10.20.563172

**Authors:** Teresa W. Lo, Han James Choi, Dean Huang, Paul A. Wiggins

**Author notes:** These two authors contributed equally (Teresa W. Lo, Han James Choi).

## Abstract

The processes of gene expression are inherently stochastic, even for essential genes required for growth. How does the cell maximize fitness in light of noise? To answer this question, we build a mathematical model to explore the trade-off between metabolic load and growth robustness. The model predicts novel principles of central dogma regulation: Optimal protein expression levels for many genes are in vast overabundance. Essential genes are transcribed above a lower limit of one message per cell cycle. Gene expression is achieved by load balancing between transcription and translation. We present evidence that each of these novel regulatory principles is observed. These results reveal that robustness and metabolic load determine the global regulatory principles that govern gene expression processes, and these principles have broad implications for cellular function.

**One-sentence summary:** Fitness maximization predicts protein overabundance, a transcriptional floor, and the balancing of transcription and translation.

## INTRODUCTION

What rationale determines the optimal transcription and translation level of a gene in the cell? Protein expression levels optimize cell fitness [1, 2]: Too low of an expression level of essential proteins slows growth by compromising the function of essential processes [3, 4], whereas the overexpression of proteins slows growth by increasing the metabolic load [5]. This trade-off naïvely predicts that the cell maximizes its fitness by a Goldilocks principle in which cells express just enough protein for function [6]; however, achieving growth robustness is nontrivial, since all processes at the cellular scale are stochastic, including gene expression [7]. This biological noise leads to significant cell-to-cell variation in protein numbers, even for essential proteins that are required for growth [8, 9]. The optimal expression program must therefore ensure robust expression of hundreds of distinct essential gene products. In this paper, we explore the consequences of growth robustness on the central dogma regulatory program.

## RESULTS

### Defining the RLTO Model

To study the consequences of growth robustness on gene expression quantitatively, we propose and analyze a minimal model: the Robustness-Load Trade-Off (RLTO) Model. The model includes three critical components: (i) Protein levels are stochastic and the single-cell growth rate depends upon them, (ii) gene transcription and translation generate a metabolic load, and (iii) cell growth is dependent on a large number of essential genes. These model characteristics result in a highly-asymmetric fitness landscape. The optimization of expression on this asymmetric landscape predicts new phenomenology absent from previous models (*e*.*g*. [10]).

The protein number *N*_*p*_ expressed from gene *i* is the product of two sequential stochastic processes: transcription and translation [11], leading to cell-to-cell variation in protein number, which we will refer to as *noise*. In our analysis, we will model gene expression using the canonical steady-state noise model [12]. (See Fig. 1A.) In this model, the numbers of proteins *N*_*p*_ for gene *i* are predicted to be gamma-distributed [13], in close agreement with observation [9]. The distribution is described by two gene-specific statistical parameters: *message number* (*μ*_*m*_), defined as the mean number of messages transcribed per cell cycle for gene *i*, and the *translation efficiency* (*ε*), the mean number of proteins translated from each message transcribed for gene *i*. The mean protein abundance is their product: *μ*_*p*_ = *μ*_*m*_*ε*. These parameters can be expressed in terms of ratios of the rates of the underlying gene expression processes, as described in Supplementary Material Sec. A 1.

**FIG. 1.**
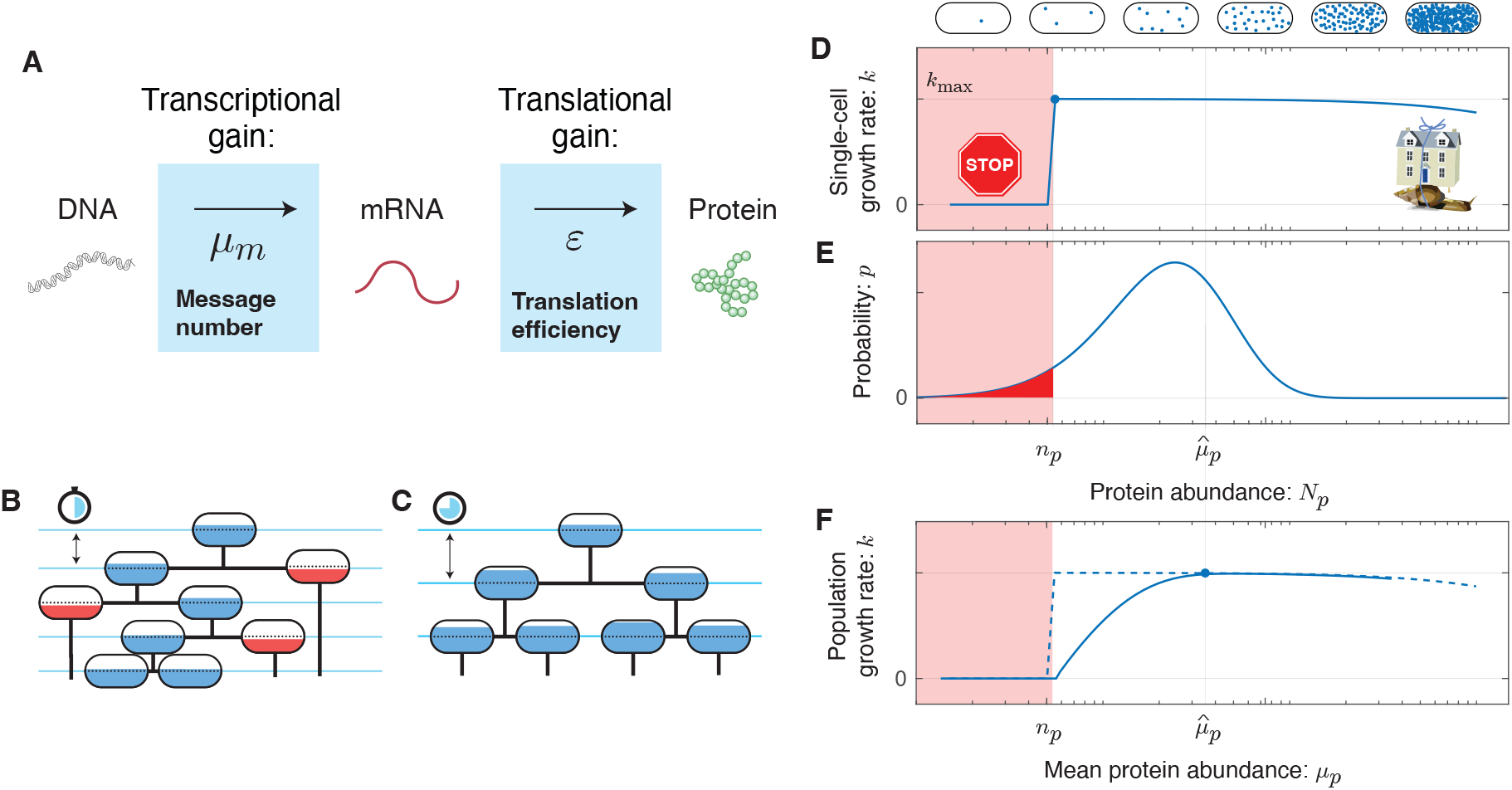
The RLTO Model. **Panel A: Gene expression is stochastic.** The central dogma describes a two-stage stochastic process where genes are first transcribed and then translated. The transcription process transcribes an average of *μ*_*m*_ messages per cell cycle. The translation process translates an average of *ε* proteins per message. **Panel B & C:** A schematic cell lineage tree is shown during exponential growth. For a specific protein *i*, the cell fill represents the protein number *N*_*p*_ relative to its threshold number *n*_*p*_ required for cell growth. **Panel B:** Reducing the mean expression level reduces doubling time; however, stochasticity in expression results in below-threshold cells (red fill) which grow slowly. **Panel C:** Increasing protein expression increases the doubling time; however, all cells are above threshold (blue fill). **Panel D: The fitness landscape as a function of protein number**. Growth arrests for protein number *N*_*p*_ smaller than the threshold level *n*_*p*_ (red) due to the failure of essential processes. High expression levels are penalized due to the metabolic cost of protein expression. This trade-off leads to a highly asymmetric fitness landscape: The relative metabolic cost of overabundance is small relative to the cost of growth arrest due to the large size of the total metabolic load *N*_0_. **Panel E: The gene expression process is stochastic**. There is significant cell-to-cell variation in protein abundance (*N*_*p*_) around the mean level (*μ* _*p*_). Even for mean expression levels significantly above the threshold level *n*_*p*_, some cells fall below threshold (red). The distribution in protein number is modeled using a gamma distribution [9], parameterized by message number *μ* _*m*_ and translation efficiency *ε*. **Panel F: The robustness load trade-off determines the optimal expression level**. The population growth rate depends on the distribution of the protein number. The asymmetry of the fitness landscape drives the optimal expression level far above the threshold level due to the high fitness cost of low protein abundance.

How should the effect of essential protein expression on growth rate be modeled in the context of the RLTO model? Much recent work has focused on cellular resource allocation to functional sectors (*e*.*g*. [14]). In this approach, an optimization is performed by the *coordinated* modulation of the abundance of all proteins in a particular sector, leading to a trade-off between functional capacities of the cell. However, in the RLTO model, the optimization is fundamentally different: We consider the *uncoordinated* modulation of the abundance of protein species *i* due to noise. For these incoherent changes, we generically expect proteins to exhibit rate-limited kinetics: Increases in the protein number *N*_*p*_ above a threshold level *n*_*p*_ has minimal effect on the rate since other chemical species (proteins, metabolites, *etc*.) are rate limiting [15]. However, if the protein number *N*_*p*_ falls below the threshold *n*_*p*_, then protein species *i* becomes rate limiting and leads to a significant slowdown of the growth rate. In the RLTO model, we coarse-grain the details of this growth slowdown as growth arrest. (See Fig. 1.) There is already some precedent for the use of this type of threshold (*e*.*g*. [16]), but we will demonstrate that the detailed form of the fitness landscape is not important. (See Supplementary Material Sec. B.) Although sufficiently detailed knowledge of the relevant molecular and cellular biology could be used to predict the protein thresholds *n*_*p*_, we will treat these as gene-specific unknown parameters.

As shown in Materials and Methods, the relative cellular fitness with respect to the expression of gene *i* can be computed by combining the fitness losses associated with robustness (Eq. 6) and metabolic load (Eq. 8):

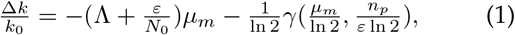

where the first term represents fitness loss due to metabolic load of transcription and translation while the second term represents loss due to the arrest of essential processes. *γ* is the regularized incomplete gamma function and the central distribution function (CDF) of the gamma distribution. (See Supplementary Material Sec. A 1 and A 2.) In summary, the model depends only on a single global parameter: the relative metabolic load Λ and three gene-specific parameters: the threshold number *n*_*p*_, the message number *μ*_*m*_ and the relative translation efficiency *ε/N*_0_. We propose that the cell is regulated to maximize the growth rate with respect to transcription (message number) and translation (translation efficiency). The fitness landscape predicted by the RLTO model for representative parameters is shown in Fig. 2A.

**FIG. 2.**
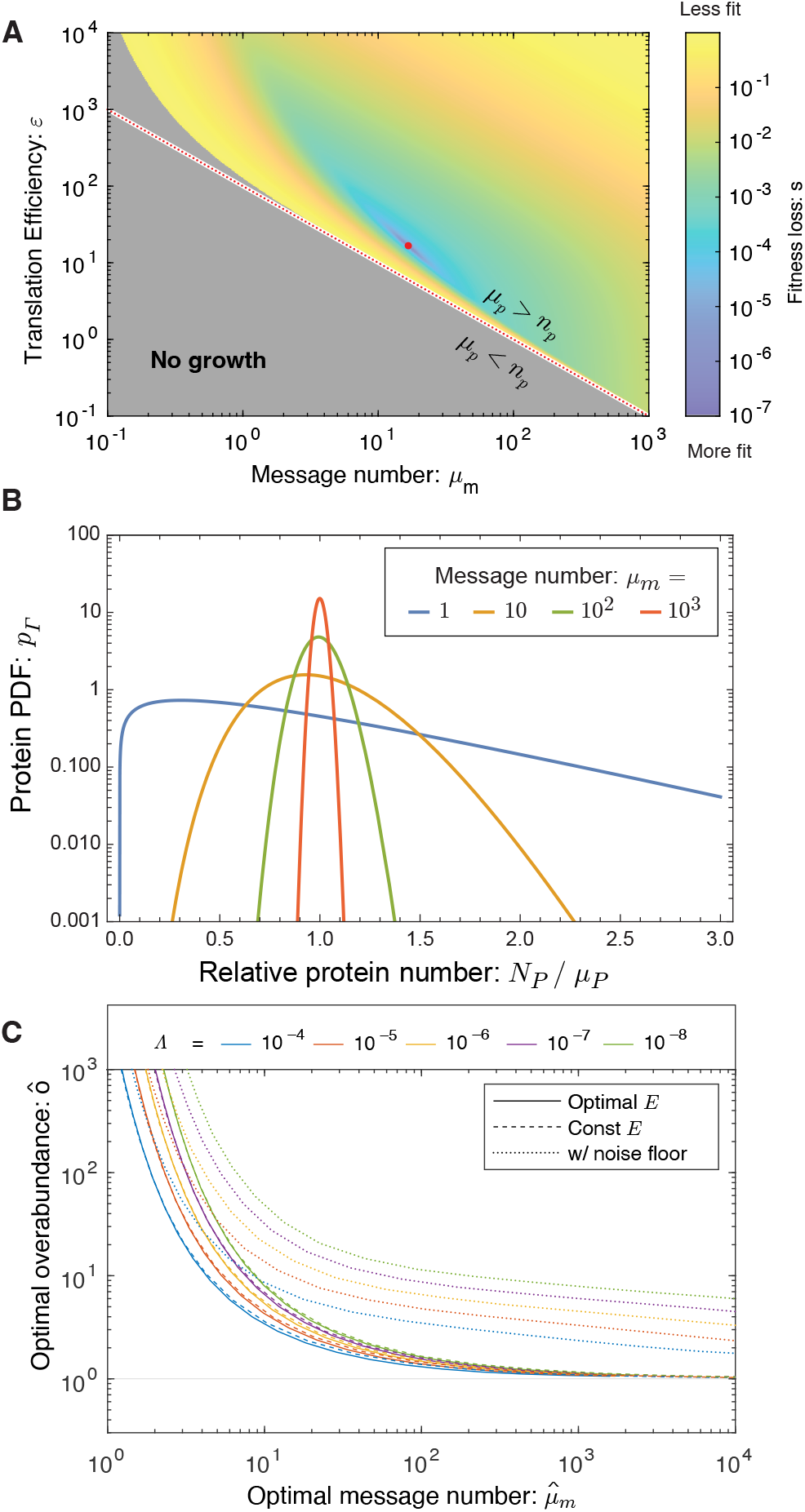
The RLTO model predicts overabundance is optimal. **Panel A: Fitness landscape determines optimal message number and translation efficiency.** The fitness loss (*s* ≡ ln *k*_max_*/k*) is shown as a function of message number (*μ* _*m*_) and translation efficiency (*ε*). The red dotted curve represents programs where the mean protein number is equal to the threshold (*μ* _*p*_ = *np*) and the red dot represents the optimal regulatory program 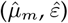. **Panel B: Gene-expression noise**. Due to the stochasticity of gene expression processes at equilibrium, the protein number *N*_*p*_ is gamma-distributed [9]. For high-expression genes, expression has low noise and the protein number is tightly distributed around its mean; however, for low-expression genes, expression is noisy and the distribution is extremely wide. **Panel C: Overabundance is optimal for all genes**. For high-expression genes, low overabundance is optimal (*μ* _*p*_ ≈ *n*_*p*_); however, for low-expression genes, vast over-abundance is optimal (*μ* _*p*_ ≫ *n*_*p*_). From a quantitative perspective, overabundance depends on the relative load Λ; however, the qualitative dependence is invariant to over an orders-of-magnitude range of values.

### RLTO predicts protein overabundance

The optimal regulatory program (*μ*_*m*_ and *ε* values) can be predicted analytically. They depend on only a single global parameter, the relative load Λ, and the gene-specific threshold number *n*_*p*_. Since the threshold number is not directly observable experimentally, we will instead predict the optimal overabundance *o*, defined as the ratio of the mean protein number to the threshold number:

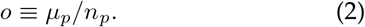

As shown in Fig. 2C, the RLTO model generically predicts that the optimal protein fraction is overabundant (*o >* 1); however, the overabundance is not uniform for all proteins. For highly-transcribed genes (*μ*_*m*_ ≫ 1) like ribosomal genes, the overabundance is predicted to be quite small (*o* ≈ 1); however, for message numbers approaching unity, the overabundance is predicted to be extremely high (*o* ≫ 1). At a quantitative level, the relation between optimal overabundance and message number depends on the relative load (Λ), but its phenomenology is qualitatively unchanged over orders of magnitude variation in Λ.

### Understanding the rationale for overabundance

To explore both the robustness of the protein overabundance prediction and to understand its mathematical rationale, we explored a collection of more complex models numerically. (Supplementary Material Sec. B.) The key mathematical feature that drives overabundance is not the assumption of growth arrest, but rather the strong asymmetry of the fitness landscape: the high cost of protein *underabundance* and the low cost of protein *overabundance*. (See Fig. 1EF.) The population growth rate (Panel 1F) can be understood qualitatively as the convolution of the single-cell growth rate (Panel 1D) with the probability density function (PDF) of the protein abundance (Panel 1E). In the RLTO model, this asymmetry is parameterized by the relative load (Λ), defined as the relative metabolic cost of transcribing an additional message. Since we estimate that Λ *<* 10^*−*5^, this cost is very low relative to the total metabolic cost of the cell, therefore we expect this asymmetry, and the prediction of the RLTO model, to be robust.

### Overabundance is observed in a range of experiments

The RLTO Model predicts that all essential proteins are overabundant. In general, the RLTO model predicts that protein numbers have very significant robustness (*i*.*e*. buffering) to protein depletion. Although this result is potentially surprising, it is in fact consistent with many studies. For instance, Belliveau *et al*. have recently analyzed the abundance of a wide range of metabolic and other essential biological processes, and conclude that protein abundance appears to be in significant excess of what is required for function [6]. Likewise, CRISPRi approaches have facilitated the characterization of essential protein depletion. The qualitative results from these experiments are consistent with over-abundance: Large-magnitude protein depletion is typically required to generate strong phenotypes [3, 17, 18]. In particular, Peters *et al*. engineered a complete collection of CRISPRi essential-gene depletion constructs in *Bacillus subtilis*. Importantly, when *dcas9* is constitutively expressed, these constructs deplete essential proteins about three-fold below their endogenous expression levels [3]; however, roughly 80% grew without measurable fitness loss in log-phase growth despite the depletion. When grouped by functional category, only ribosomal proteins were found to have statistically significant reductions in fitness [3]. As shown in Fig. 2C, the RLTO model predicts that all but the highest expression proteins are expected to show minimal fitness reductions in response to a three-fold depletion of essential enzymes. The optimality of protein overabundance explains the paradox of protein expression levels being simultaneously optimal [1] and in excess of what is required for function [3, 4, 6, 18]. Although this qualitative picture of essential protein overabundance is clear, there has yet to be a quantitative and detailed measurement of protein overabundance, and in particular, an analysis of the relationship between protein overabundance and message number.

### RLTO predicts a one-message transcription threshold

The RLTO model predicts protein overabundance, but is there a clear transcriptional signature? To analyze this question, we define the message threshold *n*_*m*_ ≡ *μ*_*m*_*/o*. (This parameterization is convenient since it is independent of the translation efficiency.) We can then analyze the relation between optimal message number and threshold message number, as shown in Fig. 3A. The model predicts that even for genes that have extremely small threshold message numbers (*e*.*g. n*_*m*_ = 10^*−*2^), the optimal message number stays above one message transcribed per cell cycle. Qualitatively, expressing messages below this level is simply too noisy even for proteins needed at the lowest expression levels. (See the blue curve in Fig. 2B corresponding to the protein number distribution of *μ*_*m*_ = 1.) The model therefore predicts a lower floor on transcription for essential genes of one message per cell cycle.

**FIG. 3.**
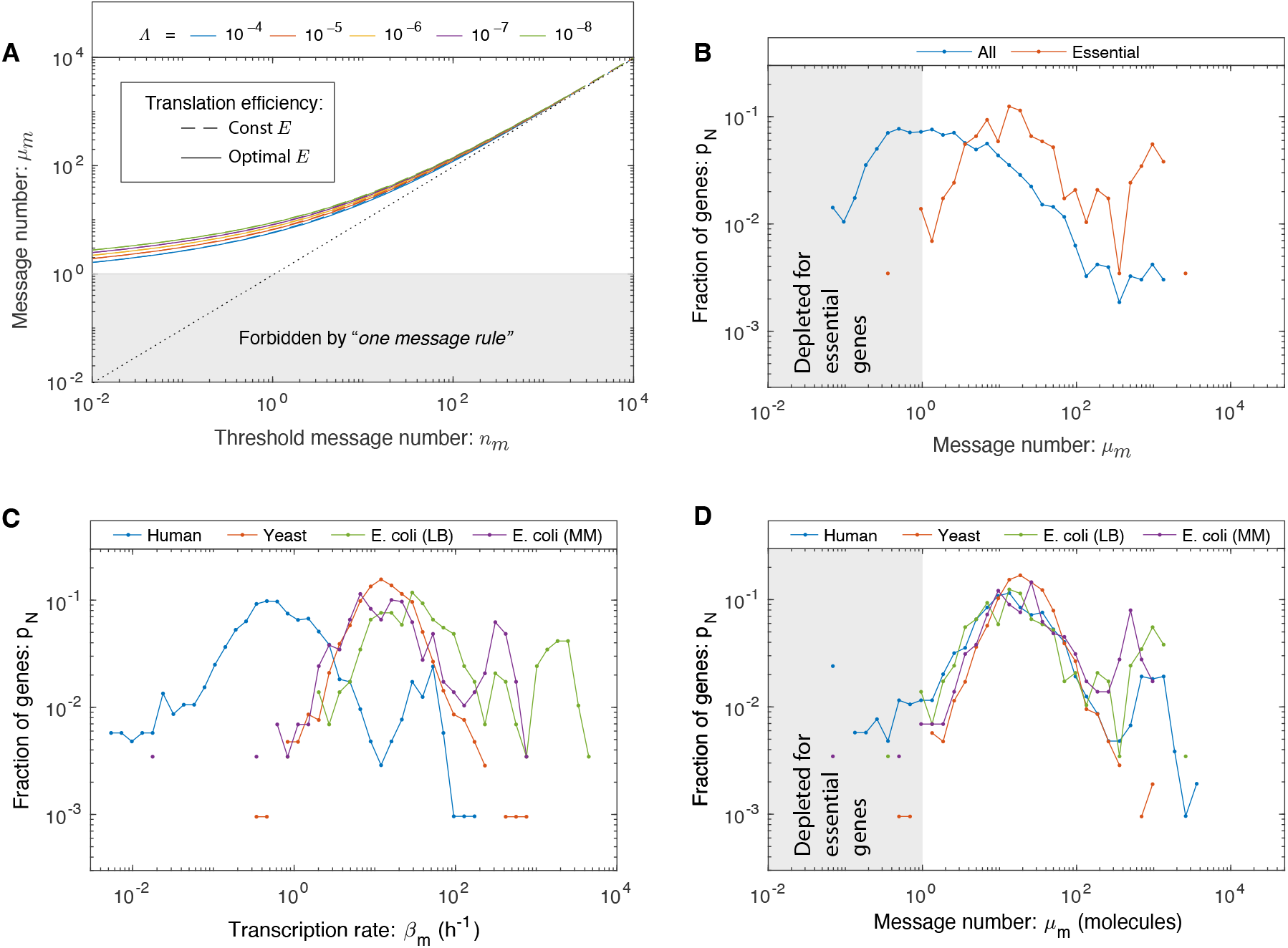
A lower threshold for transcription: The one message rule. **Panel A: RLTO predicts the one-message rule.** For high-expression genes, overabundance is low and the message number *μ* _*m*_ is predicted to be comparable to the threshold level *n*_*m*_ (dotted line); however, for low-expression genes there is a lower threshold (*μ* _*m*_ ≥ 1) below which expression is too noisy for robust growth. The threshold is weakly dependent on relative load Λ. **Panel B: A one-message threshold is observed in *E. coli* for essential genes**. A histogram shows the distribution of gene message numbers for all genes (blue) versus essential genes (orange). As predicted by the RLTO model, virtually all essential genes are expressed above the one-message-per-cell-cycle threshold. **Panel C: The distribution of transcription rates for essential genes**. No alignment is observed between the distributions of transcription rates in three evolutionarily-divergent organisms. For instance, the per gene transcription rate is significantly lower in human cells relative to *E. coli*. **Panel D: The distribution of message numbers for essential genes in three evolutionarily-divergent organisms**. The alignment of distributions of message number per gene between human, yeast, and *E. coli* (under two distinct growth conditions) reveals a nontrivial commonality between central dogma regulatory programs. We propose that the rationale for this alignment is the one-message rule that predicts that all essential genes must be expressed above one message per cell cycle. Both yeast and *E. coli* come very close to satisfying this proposed threshold; however, a greater proportion of genes in human break the one-message threshold. We speculate that this is due in part to the *ad hoc* nature of the essential-gene classification in the context of complex multicellular organisms.

### A lower threshold is observed for message number

To identify a putative transcriptional floor, we first analyzed the transcriptome in *Escherichia coli*. We hypothesize that cells must express essential genes above the one-message threshold for robust growth. The distinction between essential and nonessential genes is critical in this context, since nonessential genes can be inducibly expressed. For instance, in *E. coli*, the *lac* operon is repressed in the absence of lactose and therefore need not satisfy the one-message threshold. The transcriptional threshold is only hypothesized to apply to genes whose products are required to maintain cell fitness under the measured conditions.

We generated histograms for *E. coli* growing rapidly on rich media for these two classes of genes. (See Supplementary Material Sec. C.) The message numbers for *nonessential* genes are widely distributed, with a significant fraction of genes falling below the one-message threshold; however, only one *essential* gene is expressed below the one-message threshold (0.3% of essential genes). (See Fig. 3B.) The threshold is not sharp, but rather a smooth depletion relative to a median of 18 messages per cell cycle. This observation is consistent with the predictions of the RLTO model.

To further test this prediction, we then analyzed *E. coli* transcription under slow-growth conditions. Since these cells are less transcriptionally active, we hypothesized that this analysis would constitute a more stringent test of the one-message rule. (See Supplementary Material Sec. C.) To our surprise, although the transcription rate is indeed reduced in slow growth, the essential gene message numbers still satisfy the one-message rule (with a two gene exception, 0.7%), again consistent with the predictions of the RLTO model. (See Fig. 3D.)

Next, we analyzed eukaryotic transcriptomes in *Saccharomyces cerevisiae* (yeast) and *Homo sapiens* (human). (See Supplementary Material Sec. C.) For yeast, there is a well-defined notion of essential genes [19]. As predicted, yeast essential genes obey the one-message threshold (with two exceptions, 0.2%). (See Fig. 3D.) The interpretation is less clear-cut in human cells: An essential gene classification has been generated in the context of proliferation in cell culture [20]. In order to try to capture a generic picture, we average the human transcriptome of cell types. We find that the vast majority of essential genes obey the one-message rule; however, there are significantly more genes that break the rule (81 genes, 8%) than in the other organisms.

### Message number distribution is conserved

To what extent is this human data consistent with the RLTO model? For human cells, our test of the one-message rule is too simplistic in two respects: (i) We ignore the significant transcriptional differences associated with distinct cell types and (ii) the essential gene classification itself is defined by the ability of mutants or knock-downs to proliferate in cell culture; in marked contrast to the *in vivo* context where cell proliferation is tightly regulated [20]. Due to these subtleties, we decided to take a complementary approach: We considered the distribution of three different transcriptional statistics for each gene: transcription rate, cellular message number, defined as the average number of messages instantaneously, and message number (*μ*_*m*_), defined as the number of messages transcribed in a cell cycle. (See Supplementary Material Sec. C.) The RLTO model predicts a one-message threshold with respect to message number, but not the other two statistics. We therefore predict that the message number distributions in each organism (*E. coli*, yeast, and human) should align for low expression genes with respect to message number, but not for the other two transcriptional statistics. Consistent with the predictions of the RLTO model, there is a striking alignment of message number for essential genes between all three model organisms and growth conditions for message number. (See Fig. 3D.) This alignment is non-trivial: It is not observed with respect to other transcriptional statistics (Fig. 3C and Supplementary Material Fig. 3).

What is the significance of the similarity in the distributions of message number between organisms? Another strategy for satisfying the one message rule would be for transcription to be increased. For instance, mammalian cells have about 1000 times the number of proteins relative to bacterial cells. (See Supplemental Tab. S2.) One might therefore naively predict that the message number should be increased 1000-fold as well. This is not observed. In fact, the message number distributions of all the model organisms analyzed abut the one message threshold. The proximity to the threshold suggests that organisms do as little transcription as possible while satisfying the one message rule. This appears to be a conserved transcriptional regulatory strategy from *E. coli* to human.

### Translation efficiency is predicted to increase with transcription

What does the RLTO model predict about how the cell should balance the gene expression process between transcription and translation? Minimizing transcription (at fixed protein abundance) reduces the metabolic load; however, it decreases robustness. Growth rate maximization balances these two costs. Quantitatively, the maximization of the growth rate (Eq. 1) with respect to the translation efficiency can be performed analytically, predicting the optimal translation efficiency, shown in Fig. 4A. We provide an exact expression in the Supplementary Material Sec. A 4; however, an approximate expression for the translation efficiency is more clearly interpretable:

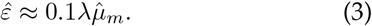

**FIG. 4.**
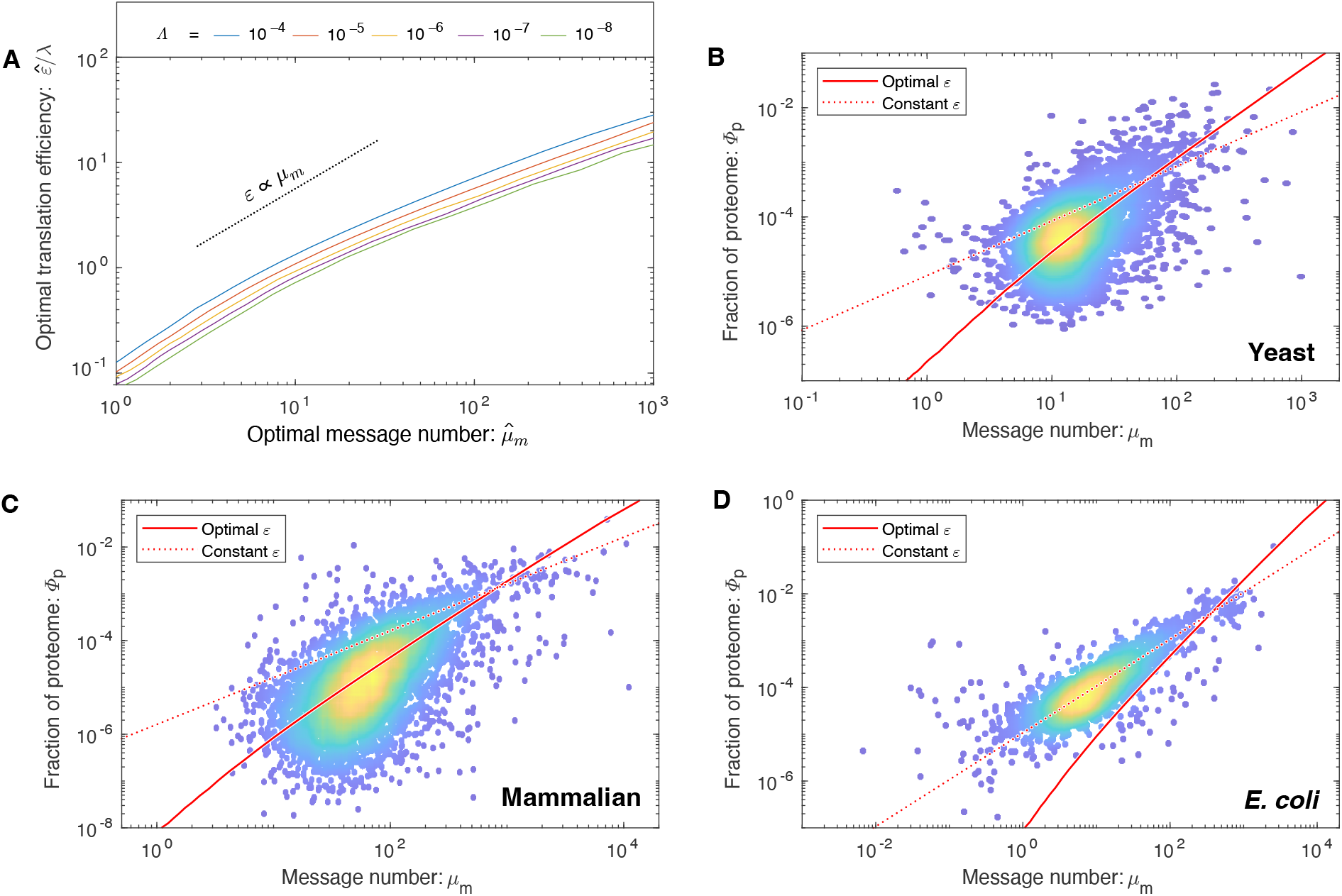
How are transcription and translation balanced? **Panel A: The RLTO model predicts load balancing.** The ratio of the optimal translation efficiency 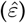 to the message cost (*λ*) is roughly independent of the relative load (Λ). The translation efficiency *ε* is predicted to be roughly proportional message number *μ* _*m*_. **Panel B: RLTO predicts the protein-message-abundance relation in yeast**. The observed proteome fraction is compared to two models: the RLTO optimal model (solid red line) and constant-translation-efficiency model (dotted red line). Both models make parameter-free predictions. The RLTO optimum predicts the global trend. (Data from Ref. [21].) **Panel C: Mammalian proteome fraction**. The RLTO prediction (solid) is superior to the constant-translation-efficiency prediction (dashed). **Panel D: *E. coli* proteome fraction**. In contrast, the constant-translation-efficiency prediction (dashed) is superior to RLTO prediction (solid).

The optimal translation efficiency has two important qualitative features for central dogma regulation. The first prediction is that as the message cost (*λ*) rises, the optimal translation efficiency 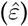 increases in proportion while the message number decreases. We present evidence for this prediction in the Supplementary Material Sec. A 11.

The second prediction is that the optimal translation efficiency is also approximately proportional to message number 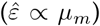. Therefore, the RLTO model predicts that low expression levels should be achieved with low levels of transcription and translation, whereas high-expression genes are achieved with high levels of both.

We call this relation between optimal transcription and translation the *load balancing principle*. The most direct test of load balancing is measuring the protein-message abundance relation. Due to load balancing, the RLTO model predicts protein number (and proteome fraction) to scale like:

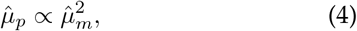

whereas a constant-translation-efficiency model has linear scaling (*μ*_*p*_ ∝ *μ*_*m*_). Computing proteome fraction, rather than protein number, results in a parameter-free prediction. (See Supplemental Sec. A 12.)

### Load balancing is observed in eukaryotic cells

To test the RLTO predictions, we compare observed proteome measurements in three evolutionarily divergent species, *E. coli* [22], yeast [21] and mammalian cells [23], to two models: the RLTO and the constant-translation-efficiency models. The results of the parameter-free predictions are shown in Fig. 4BCD for each organism. The RLTO model clearly captures the global trend in the proteome-fraction message-number relation in eukaryotic cells and a direct fit to a power law with an unknown exponent is consistent with Eq. 4 (Supplementary Material Sec. D 4 c).

In *E. coli*, the constant-translation-efficiency model better describes the data. Why does this organism appear not to load balance? In the supplementary material, we demonstrate that the observed translation efficiency is consistent with the RLTO model, augmented by a ribosome-per-message limit. Hausser *et al*. have proposed just such a limit, based on the ribosome footprint on mRNA molecules [10]. (See Supplementary Material Sec. A 17.) Although this augmented model is consistent with central dogma regulation in *E. coli*, it is not a complete rationale. This proposed translation-rate limit could be circumvented by increasing the lifetime of *E. coli* messages, which would increase the translation efficiency. A more in-depth analysis specific to *E. coli* is needed to understand why the observed message life-time is so short.

### RLTO model predicts observed noise in yeast

Although the protein fraction measurements support the RLTO predictions for the translation efficiency in eukaryotic cells, these measurements do not provide a compelling rationale for why load balancing maximizes the growth rate. To understand its rationale, we explore its implications for noise.

In a typical biological context, *μ*_*m*_ ≪ *ε* and as a result, noise production is dominated by the transcription step of the gene expression process [12, 13]. (A table of central dogma parameters for each model organism appears in the Supplementary Material Tab. S2.) Quantitatively, the canonical steady-state noise model predicts that the noise should be inversely related to the message number [12, 13]:

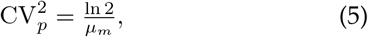

however, it is the relation between mean protein abundance *μ*_*p*_ and noise 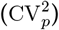 which is typically reported [8, 9]. Based on the scaling of the optimal translation efficiency with the message number in eukaryotic cells (Eq. 3), we find the protein number to scale with message number (Eq. 4), which predicts that noise should scale with protein abundance 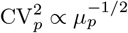 in yeast (see Supplementary Material Sec. D 1); however, due to the observed absence of translation-efficiency scaling in bacteria, the noise should scale as 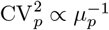 in bacteria, as observed [9]. Does the yeast noise show the predicted scaling? The parameter-free RLTO noise prediction closely matches the observed noise in both magnitude and scaling, as shown in Fig. 5.

**FIG. 5.**
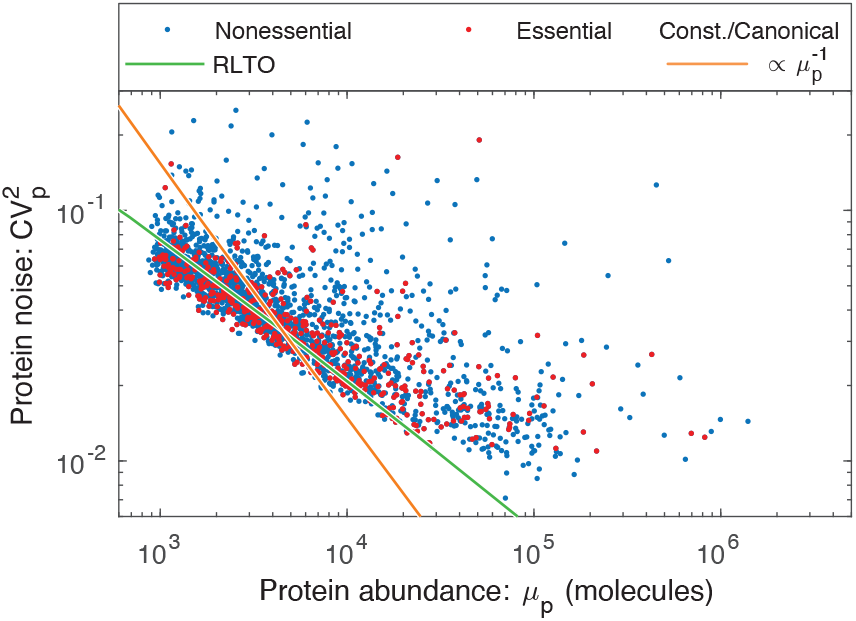
RLTO predicts the magnitude of noise in yeast. The observed gene expression noise is yeast is shown for essential and nonessential genes. Two protein-message abundance models are compared to the data: The RLTO model (green) versus the constant-translation-efficiency (canonical model, orange). The RLTO model predicts both the magnitude of the noise, as well as its scaling with protein abundance. The reduced slope of the RLTO model is the consequence of load balancing, which reduces the noise for the noisiest, low-expression genes. (Data from Ref. [8].)

### Reducing noise is the rationale for load balancing

This noise analysis also provides a conceptual insight into the rationale for load balancing. The load balanced (RLTO-green) and constant-translation-efficiency (orange) predictions for the noise are shown in Fig. 5. Load balancing results in decreased noise for low-expression, noisy genes over what is achieved with constant translation efficiency. This decreased noise is predicted to increase growth robustness. In principle, the noise could be reduced further by tipping the balance even more towards transcription; however, the RLTO model predicts that this approach is too metabolically costly, and the optimal strategy is that observed for noise scaling in yeast.

## DISCUSSION

### What are the biological implications of noise?

Many important proposals have been made, including bet-hedging strategies, the necessity of feedback in gene regulatory networks, *etc*. [7]. Our model suggests that overcoming cell-to-cell variation may fundamentally reshape the metabolic budget: Typically, proteins constitute 50-60% of the dry mass of the cell [5] and therefore overabundance could increase the overall protein budget by a significant factor. Why does the cell tolerate this significant increase in metabolic load above what would be predicted by a resource allocation analysis (*e*.*g*. [24])? This strategy dramatically reduces the consequence of stochastic expression of proteins on the rate of single-cell proliferation.

A second source of stochasticity, environmental fluctuations, has been proposed as a rationale for overabundance [25], especially in the context of metabolic genes [26]. In short, cells express protein to hedge against starvation [25] or changes in the carbon source, *etc*. [26].

How does this hypothesis compare to our growth robustness hypothesis? There are some similarities between these environmental-fluctuation models and the RLTO model: In both models, it is a fluctuations-based mechanism that drives overabundance; however, there are important distinctions between the model predictions. In the environmental fluctuation model, there is a trade-off between log-phase fitness and the rapidity of adaptation [25]; whereas in the RLTO model, over-abundance corresponds to the log-phase optimum. Organisms experiencing prolonged periods of balanced growth would therefore be expected to reduce over-abundance. Furthermore, the environmental fluctuation model most naturally explains overabundance for proteins related to metabolic processes, whereas the RLTO model predicts overabundance generically, dependent only on message number, which appears to be much more consistent with experiments exploring essential-protein depletion [4].

### Implications for nonessential genes

In our analysis, we have focused on essential genes in order to motivate the growth-threshold in the RLTO model. To what extent do nonessential genes share the same optimization? In support of the proposal that RLTO optima describe nonessential genes is the success of the model in predicting the translation efficiency for all genes, not just essential genes. (See Fig. 4 and Fig. 5.) Furthermore, the definition of a gene as *essential* depends on context: For instance, in the context of *E. coli* growth on lactose, the gene *lacZ* is essential, although it is nonessential on other carbon sources [27]. Under growth conditions where the *lacZ* gene is essential, we predict that LacZ should be overabundant, consistent with observation [26]. Finally, our modeling suggested that RLTO model phenomenology is the results of asymmetry of the cost of under versus overabundance. For nonessential genes whose activity significantly increases fitness, we still expect fitness asymmetry due to the low relative metabolic cost of increased expression. We therefore expect all gene products, most especially those with low expression, to be overabundant, under conditions where their activity increases fitness.

### Implications of overabundance for inhibitors

The generic nature of overabundance, especially for low-expression proteins, has important potential implications for the targeting of these proteins with small-molecule inhibitors (*e*.*g*. drugs). For the highest expression proteins, like the constituents of the ribosome, relatively small decreases in the active fraction (*e*.*g*. a threefold reduction) are expected to lead to growth arrest [3]. This may help explain why inhibitors targeting translation make such effective antimicrobial drugs. However, we predict that the lowest expression proteins require a much higher fraction of the protein to be inactivated, with the lowest-expression proteins expected to need more than a 100-fold depletion. This predicted robustness makes these proteins much less attractive drug targets [28].

### The principles that govern central dogma regulation

We propose that robustness to noise fundamentally shapes the central dogma regulatory program for all genes and predicts a number of key regulatory principles. (See Fig. 6.) For high-expression genes, load balancing implies that gene expression consists of both high-amplification translation and transcription. The resulting expression level has low overabundance relative to the threshold required for function. In contrast, for essential low-expression genes, a three-fold strategy is implemented: (i) overabundance raises the mean protein levels far above the threshold required for function, (ii) load balancing, and (iii) the one-message rule ensures that message number is sufficiently large to lower the noise of inherently-noisy, low-expression genes. We anticipate that these regulatory principles, in particular protein overabundance, will have important implications, not only for our understanding of central dogma regulation specifically, but for understanding the rationale for protein expression level and function in many biological processes.

**FIG. 6.**
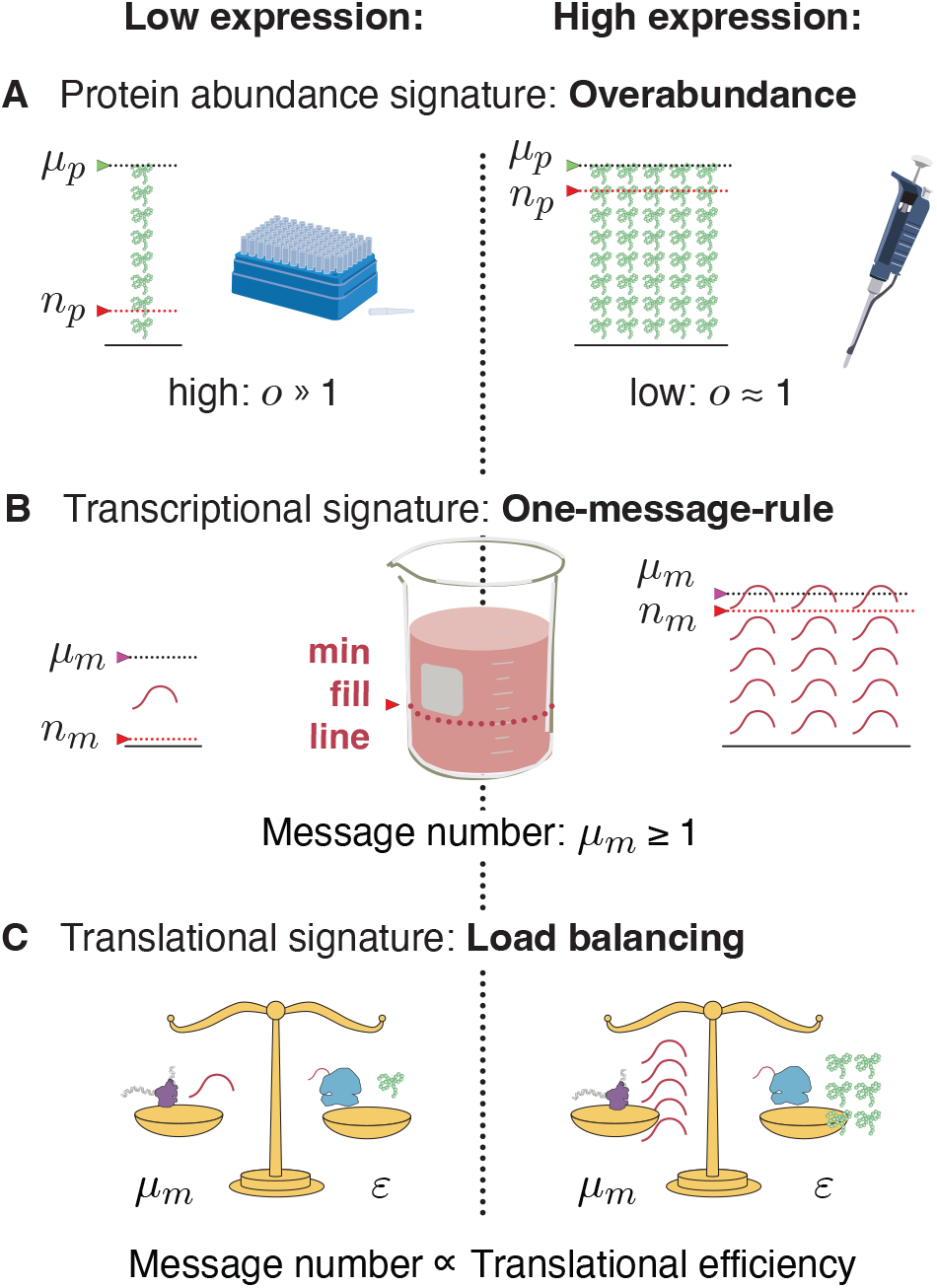
Central dogma regulatory principles. **Panel A: Over-abundance.** Low-expression essential genes are expressed with high overabundance; whereas, high-expression essential genes are expressed with low overabundance. Lab supply analogy: Low-cost items that are used stochastically (*e*.*g*. pipette tips) are purchased in great excess, while the higher cost items that are less stochastic (*e*.*g*. pipette) are purchased as needed. **Panel B: One-message rule**. Robust expression of essential genes requires them to be transcribed above a threshold of one message per cell cycle. **Panel C: Load balancing**. In eukaryotic cells, optimal fitness is achieved by balancing transcription and translation: The optimal message number is proportional to the optimal translation efficiency. High (low) expression levels are achieved by high (low) levels of transcription followed by high (low) levels of translation per message.

## MATERIALS AND METHODS

### RLTO model

The effect of stochastic cell arrest can be implemented analytically as follows: The probability of growth is the probability that all essential proteins are above threshold, *P*_+_. The population growth rate *k* is [29]:

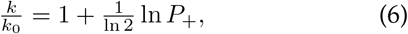

for a population of cells subject to stochastic arrest with probability 1 *− P*_+_ per cell cycle where *k*_0_ is the growth rate of the non-arrested cells. For each gene *i*, the canonical steady-state noise model predicts the protein number CDF in terms of message number *μ*_*m*_ and translation efficiency *ε* [12]. Assuming the below-threshold probability is small, the probability that the cell is below threshold for gene *i* is:

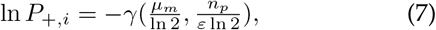

where *γ* is the regularized incomplete gamma function and the CDF of the gamma distribution. (See Supplementary Material Sec. A 2.)

While protein underabundance slows cell growth by the arrest of essential processes, protein overabundance slows growth by increasing the metabolic load. To implement the metabolic-load contribution to cell fitness, we use a minimal model that realizes the metabolic cost of both transcription and translation that is analogous to those previously used in the context of resource allocation (*e*.*g*. [14]). The metabolic load of transcription and translation of gene *i* is:

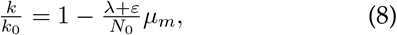

where *k*_0_ is the growth rate in the absence of the metabolic load of gene *i, N*_0_ is the total cellular metabolic load, and *λ* is the metabolic message cost. (See the Supplementary Material Sec. A 2 for a detailed development of the model.) The *λ*-term represents the metabolic cost of transcription and the *ε*-term represents the metabolic cost of translation of gene *i*. We define the relative load as Λ ≡ *λ/N*_0_ as the ratio of the metabolic load of a single message to the total metabolic cost of the cell. In *E. coli*, we estimate that Λ is roughly 10^*−*5^ and it is smaller still for eukaryotic cells.

## Data analysis

We provide a detailed description of the data analysis for the one message rule, loading balancing, and the noise analysis in the Supplementary Material.

## Acknowledgments

The authors would like to thank B. Traxler, A. Nourmohammad, J. Mougous, K. Cutler, M. Cosentino-Lagomarsino, S. van Teeffelen, C. Manoil, L. Gallagher, J. Bailey and S. Murray.

## Funding

National Institutes of Health grant R01-GM128191

## Author contributions

T.W.L., H.J.C., D.H. and P.A.W. conceived the research, performed the analysis, and wrote the paper.

## Competing interests

The authors declare no competing interests.

## Data availability

All data needed to evaluate the conclusions in the paper are present in the paper and/or the Supplementary Materials.

## Supplementary Materials

Supplementary Text

Figs. S1 to S12

Tables S1 to S3

Data Tables S1 to S11

## Supplementary Material

## Appendix A Detailed development of the RLTO model

In this section, we provide a detailed development of the RLTO model. First, we describe the stochastic kinetic model for the central dogma, which introduces key quantities for the RLTO model (Sec. A 1). Next, we provide a derivation of the growth rate as a function of the model parameters (Sec. A 2) as well as other methods (Secs. A 4, A 12, A 15, and A 18). For each of the results discussed in the main paper, we provide more detailed analyses, which include both supplemental results (Secs. A 8, A 9, A 11, A 13, and A 16) that support the story described in the main paper, as well as supplemental discussions (Secs. A 3, A 5, A 6, A 7, A 14, and A 17).

### 1. Methods: Detailed description of the noise model

#### a. Stochastic kinetic model for the central dogma

The canonical steady-state noise model for the central dogma describes multiple steps in the gene expression process [9, 12, 13]: Transcription generates mRNA messages. These messages are then translated to synthesize the protein gene products [11]. Both mRNA and protein are subject to degradation and dilution [30]. At the single cell level, each of these processes are stochastic. We will model these processes with the stochastic kinetic scheme [11]:

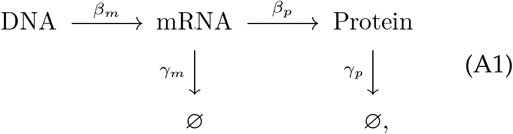

where *β*_*m*_ is the transcription rate (s^*−* 1^), *β*_*p*_ is the translation rate (s^*−*1^), *γ*_*m*_ is the message degradation rate (s^*−*1^), and *γ*_*p*_ is the protein effective degradation rate (s^*−*1^). The message lifetime is 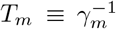. For most proteins in the context of rapid growth, dilution is the dominant mechanism of protein depletion and therefore *γ*_*p*_ is approximately the growth rate [9, 31, 32]: *γ*_*p*_ = *T* ^*−*1^ ln 2, where *T* is the doubling time.

#### b. Statistical model for protein abundance

To study the stochastic dynamics of gene expression, we used a stochastic Gillespie simulation [33, 34]. (See Sec. A 1 c.) In particular, we were interested in the explicit relation between the kinetic parameters (*β*_*m*_, *γ*_*m*_, *β*_*p*_, *γ*_*p*_) and experimental observables.

Consistent with previous reports [12, 13], we find that the distribution of protein number per cell (at cell birth) was described by a gamma distribution: *N*_*p*_ ∼ Γ(*a, θ*), where *N*_*p*_ is the protein number at cell birth and Γ is the gamma distribution which is parameterized by a scale parameter *θ* and a shape parameter *a*. (See Sec. A 1 d.) We refer to this distribution as the *canonical steady-state noise model*; The relation between the four kinetic parameters and these two statistical parameters has already been reported, and have clear biological interpretations [13]: The scale parameter:

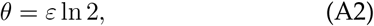

is proportional to the translation efficiency:

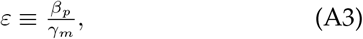

where *β*_*p*_ is the translation rate and *γ*_*m*_ is the message degradation rate. *ε* is understood as the mean number of proteins translated from each message transcribed. The shape parameter *a* can also be expressed in terms of the kinetic parameters [13]:

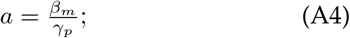

however, we will find it more convenient to express the scale parameter in terms of the cell-cycle message number:

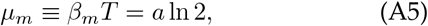

which can be interpreted as the mean number of messages transcribed per cell cycle. Forthwith, we will abbreviate this quantity *message number* in the interest of brevity.

**FIG. S1.**
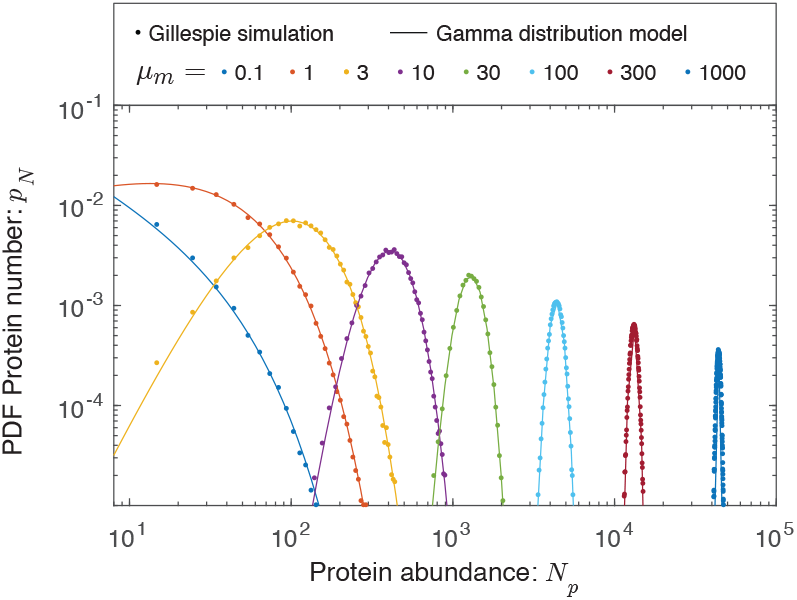
The protein abundance is approximately gamma distributed. Protein abundance was modeled for eight different transcription rates using a Gillespie simulation, including the stochastic partitioning of the proteins between daughter cells at cell division. The range in abundance matches the observed range of expression levels in the cell. We observed that the simulated protein abundances were well fit by gamma distributions.

#### c. Gillespie simulation of stochastic kinetic scheme

Protein distributions based on the kinetic scheme defined in Sec. A 1 a were simulated with a Gillespie algorithm, with specific parameter values for *E. coli*. Assuming the lifetime of the cell cycle (*T* = 30 min) [35], mRNA lifetime (*T*_*m*_ = 2.5 min) [36], and translation rate (*β*_*p*_ ≈ 500 hr^*−*1^), the protein distributions for several mean expression levels were numerically generated for exponential growth with 100,000 stochastic cell divisions, with protein partitioned at division following the binomial distribution.

The gamma distributions for each mean message number with scale and shape parameters determined by the corresponding translation efficiency and message number 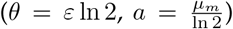 as used for the Gillespie simulation were also plotted with the protein distributions. We observe an excellent match between these Gillespie simulations and the canonical steady-state statistical noise model (*i*.*e*. gamma function) as shown in Fig. S1.

#### d. Gamma function and distribution conventions

There are a number of conflicting conventions for the gamma function and distribution arguments. We will use those defined on Wikipedia and the CRC Encyclopedia of Mathematics [37]. The gamma distributed random variable *X* will be written:

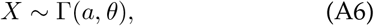

where *a* is the shape parameter and *θ* is the scale parameter. The PDF of the distribution is:

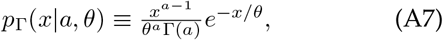

where Γ(*a*) is the gamma function. The CDF is therefore:

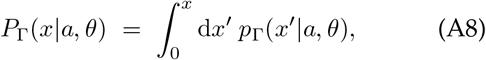

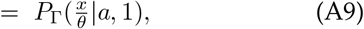

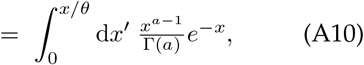

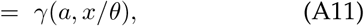

where *γ* is the regularized incomplete gamma function.

### 2. Methods: The derivation of the RLTO growth rate

#### a. The metabolic load of protein and the resource allocation model

To model the effect of the metabolic load on cell growth, we will expand on a model used by Hwa and co-workers [14]. For conciseness, we will call the original model the *resource allocation model*.

Consider a cell where the total number of proteins in the cell is *N*. The synthesis of these proteins requires two sets of processes: (i) the metabolic processes responsible for synthesizing the precursors (*i*.*e*. amino acids, *etc*) and (ii) the translation process. Proteins involved in the metabolic processes be referred to as the *P* sector and number *N*_*P*_. The proteins involved in the translational process will be referred to as the *R* sector and number *N*_*R*_. In addition to the P and R sectors, there is a third Q sector with protein number *N*_*Q*_. The total protein number per cell is therefore:

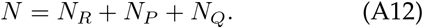

The proteome fractions are defined Φ_*X*_≡ *N*_*X*_*/N* and have the normalization condition:

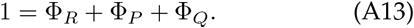

The key assumption in the resource allocation model is that the abundances of the R and P sectors can change in size to accommodate changes in the nutrient quality and translation load associated with a particular growth condition [14]. In contrast, the Q sector has a fixed proteome fraction, Φ_*Q*_, irrespective of growth conditions. In the resource allocation model, the size of these adjustable R and P sectors are chosen to optimize the growth rate *k*.

The condition for balanced growth requires that the overall protein output of the translation process match the growth rate:

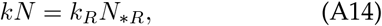

where *k*_*R*_ is the effective translation rate per protein and *N*_*\*R*_ is the number of productive R sector proteins, which is subset of the total number *N*_*R*_:

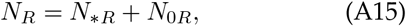

and *N*_0*R*_ represents unproductive R sector protein. We can rewrite Eq. A14 in terms of the proteome fraction:

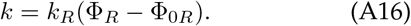

In the expression above, we will assume that the parameters *k*_*R*_ and Φ_0*R*_ are fixed, but the total fraction Φ_*R*_ is chosen to optimize the growth rate *k*.

For any productive sectors *i*, we will write analogous equations to Eq. A16 linking sector fraction size Φ_*i*_ to function:

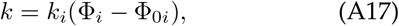

where, as before, Φ_0*i*_ represents a fixed-size fraction of unproductive protein.

To determine the unknown optimum growth rate and sector sizes, Eq. A17 can be re-written:

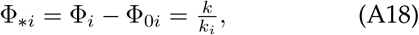

and then summed over all sectors (excluding Q):

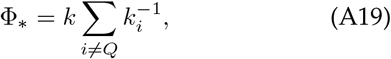

where we define the total productive fraction of the proteome:

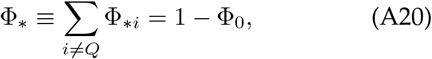

and Φ_0_ represents the total fraction of unproductive protein:

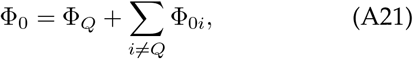

including the entire Q sector. Since Φ_*Q*_ and the Φ_0*i*_ are all assumed to be fixed, Eq. A19 determines the growth rate. (In *E. coli*, Hwa and coworkers estimate that Φ_*\**_ ≈ 0.55.)

To understand the meaning of Eq. A19, we first define an ideal growth rate as

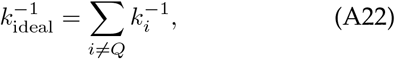

which would be the growth rate in the absence of unproductive protein; however, due to the presence of the unproductive protein, the growth rate is proportional to the productive fraction:

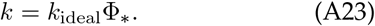

The optimal protein fractions can be determined using Eq. A18:

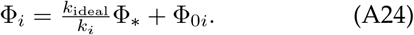

How does the growth rate change when the unproductive protein fraction is changed by *δ*Φ? The productive fraction is reduced:

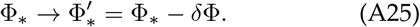

The ratio of the new growth rate *k*^*′*^ to the original is therefore:

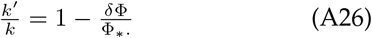

Note that our generalized resource allocation model is written for arbitrary number of functional sectors *i* ≠ *Q* and the key determinant of the change in growth rate is the fraction of productive protein Φ_*\**_. This equation will be used to model the fitness cost of the metabolic load.

#### b. The metabolic load of mRNA

What is the cost of transcription? It is perhaps useful to first consider the estimates of biosynthetic cost of macromolecules in the cell in descending order in *E. coli* [38]:

| Macromolecule | Biosynthetic cost<br>($10^9$ ATP) |
| --- | --- |
| Protein | 4.5 |
| Phospholipid | 3.2 |
| RNA | 1.6 |
| Lipopolysacchride | 3.8 |
| DNA | 0.35 |
| Peptidoglycan | 0.17 |
| Glycogen | 0.03 |

So clearly the cost of RNA is itself not insignificant. Although a significant fraction of the RNA is rRNA rather than mRNA, the mRNA itself in *E. coli* undergoes multiple rounds of transcription due to its short lifetime, increasing its cost to what is required to synthesize the molecules observed in the *E. coli* cell at any time *t*. Furthermore, transcription is dependent on protein enzymes, which themselves must be synthesized. We therefore conclude from this estimate that the cost of transcription is likely a significant determinant of the metabolic load.

Experimentally, Kafri and coworkers have measured the fitness cost of transcription and translation independently using the DAmP (Decreased Abundance by mRNA Perturbation) system in *Saccharomyces cerevisiae* [39]. As expected, they report that the metabolic cost of transcription is comparable to translation and that the reduction in growth rate is linear in transcription, in close analogy to Eq. A26.

#### c. Metabolic load in the RLTO model

To produce a minimal model to study the trade-off between robustness and metabolic load, we must consider both the metabolic cost of transcription and translation. We will write that the metabolic load (in protein equivalents) associated with gene *i* is:

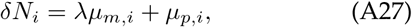

where *λ* is the message cost, the metabolic load associated with an mRNA molecule relative to a single protein molecule of the gene product. *μ*_*m,i*_ is the mean number of messages transcribed per cell cycle (mRNA molecules per cell cycle) for gene *i. μ*_*p,i*_ is the mean number of protein translated per cell cycle for gene *i*. We will describe the mean protein number in terms of the translation efficiency *ε*_*i*_, the number of proteins translated per message:

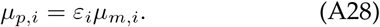

How does the cell growth rate change due to the metabolic load associated with the expression of gene *i*? The change in the metabolic load is:

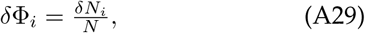

where *N* represents the total metabolic load of all components of the cell, in units of protein equivalents. Using Eq. A26, the resulting change in growth rate is:

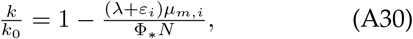

where *k*_0_ is the growth rate in the absence of the metabolic load of gene *i*.

In our analysis, the exact size of the total metabolic load *N* will not be important. In the interest of simplicity we will therefore adsorb the productive fraction Φ_*\**_ into an effective total metabolic load:

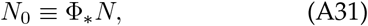

and write a concise relation between the load from gene *i* and the growth rate:

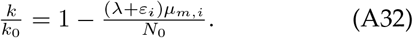

This equation has an intuitive interpretation: growth slows in proportion to the relative added metabolic load. Since Φ_*\**_ is order unity, we will ignore the distinction between the *N* and *N*_0_ quantities hence forth.

Although the global parameters *N*_0_ and *λ* provide an intuitive representation of the model, the relative growth rate depends on fewer parameters. Let *k* and *k*_0_ be the growth rates in the presence and absence of the metabolic load of gene *i*. The relative growth rate is:

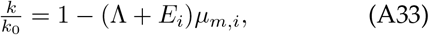

where we have introduced two new reduced parameters: the relative load, defined as Λ ≡ *λ/N*_0_, represents the ratio of the metabolic load of a single message to the total load and the relative translation efficiency, defined *E*_*i*_≡ *ε*_*i*_*/N*_0_, which is the ratio of the number of proteins translated per message to the total metabolic load *N*_0_. (Note that due to the high multiplicity *N*_0_ ≫ (*λ* + *ε*_*i*_)*μ*_*m,i*_, we can ignore the distinction between *N*_0_ and 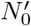 in the denominator.) If we neglect the difference between the total metabolic load and the number of proteins, the proteome fraction for gene *i* is:

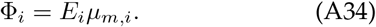

Both reduced parameters, Λ and *E*_*i*_ are extremely small. In *E. coli*, we estimate that both Λ and *E*_*i*_ are roughly 10^*−*5^ and they are smaller still for eukaryotic cells. (See Sec. A 18.)

#### d. Growth rate with stochastic arrest

For completeness, we provide a derivation of the growth rate with stochastic cell-cycle arrest that we have previously described [29]. Starting from the exponential mean expression for the population growth rate [29]:

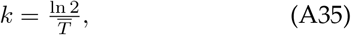

where *k* is the population growth rate and

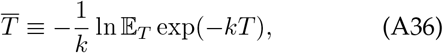

is the exponential mean, where *T* is the stochastic cell cycle duration and E is the expectation operator [29, 40]. We take a coarse-grained model which considers changes in growth rate due to fluctuations in protein number to be negligible. Note that Eq. A36 is equivalent to the *Euler-Lotka* equation [41, 42].

Let *P*_+_ be the probability of growth. When the cells are growing, the cell cycle duration *τ* is determined by the metabolic load predictions (Eq. A32). The probability mass function is therefore:

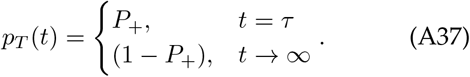

Evaluating the expectation in Eq. A36 gives:

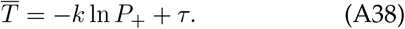

Using Eq. A35, we can solve for the growth rate *k*:

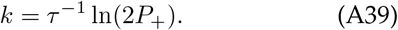

As expected, the growth rate goes down as the probability of growth *P*_+_ decreases, stopping completely at 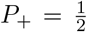. We can then compute the ratio of the growth with (*k*) and without arrest (*k*_0_):

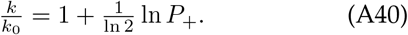

where *k*_0_ is computed by evaluating Eq. A39 at *P*_+_ = 1.

#### e. RLTO growth rate

In the RLTO model, we will assume the probability of growth is the probability that all essential protein numbers are above threshold. We will further assume that each protein number is independent, and therefore:

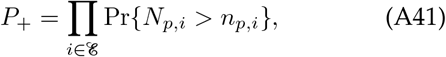

where *E* is the set of essential genes. Clearly, this assumption of independence fails in the context of polycistronic messages. We will discuss the significance of this feature of bacterial cells elsewhere, but we will ignore it in the current context.

As we will discuss, the probability of arrest of any protein *i* to be above threshold is extremely small. It is therefore convenient to work in terms of the CDFs which are very close to zero:

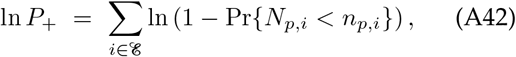

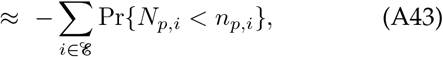

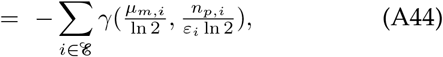

where *γ* is the regularized incomplete gamma function and the CDF of the gamma distribution (see A 1 b and A 1 d).

**FIG. S2.**
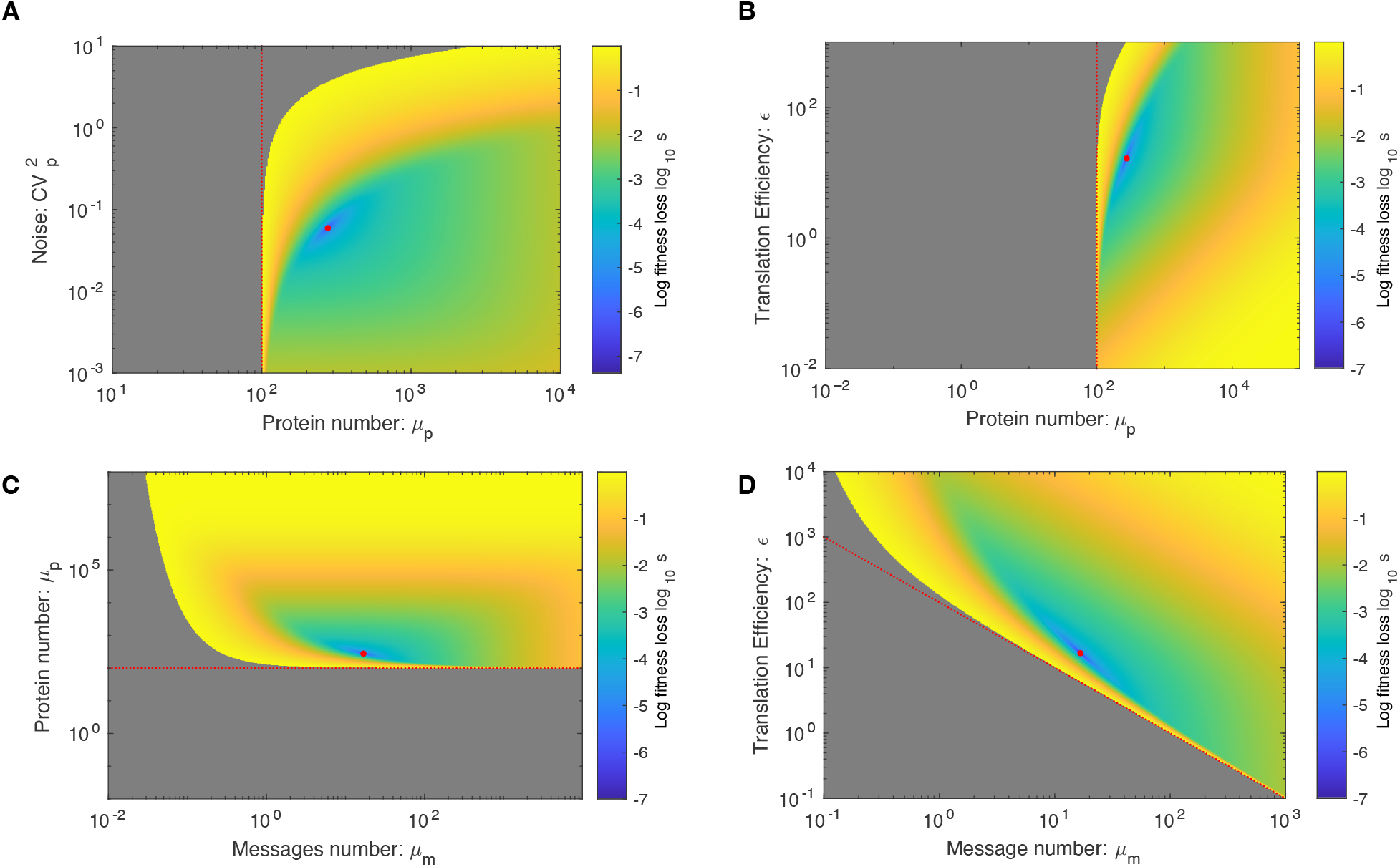
Four perspectives on the fitness landscape. In each landscape, the red circle represents the fitness optimum. The red dotted line represents the mean protein number equal to the protein threshold *n*_*p*_ = 10^2^. Here, fitness is quantified by the log growth rate: *s* = ln *k*_max_*/k*. **Panel A: Mean protein number versus noise. Panel B: Mean protein number versus translation efficiency. Panel C: Mean message versus protein numbers. Panel D: Message number versus translation efficiency**.

#### f. When the ln approximation is avoided…

The approximation discussed in the previous section is extremely well justified at the optimal central dogma parameters; however, there are a set of figures where we cannot use it. In the fitness landscape figures (Fig. S2), we compute the fitness not just at the optimal values but far from them. When cell arrest has a large effect on the growth rate, we cannot approximate the natural log with a series expansion, and we must use the full expression in Eq. A42.

#### g. Single-gene equation

By summing the fitness losses from the metabolic load and cell arrest (Eqs. A32, A40, and A44), we can write an expression for the growth rate including contributions from essential gene *i*:

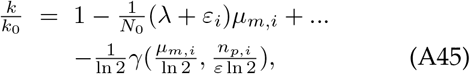

where the second term on the RHS represents the fitness loss due to the metabolic load and the third term represents the fitness loss due to stochastic cell arrest due to protein *i* falling below threshold. The fitness landscape for different gene expression parameters is shown from four different perspectives in Fig. S2. From this point forward, we will drop the subscript *i* for the sake of brevity unless otherwise noted.

#### h. Summary of RLTO parameter values for figures

The parameter values for the RLTO model used for each figure in the paper are shown in Tab. S1.

### 3. Discussion: The fitness landscape of a trade-off

The fitness landscape predicted by the RLTO model for representative parameters is shown in Fig. S2. The figure displays a number of important model phenomena: There is no growth for mean protein number *μ*_*p*_ below the threshold number *n*_*p*_, and for high noise, *μ*_*p*_ must be in significant excess of *n*_*p*_. Rapid growth can be achieved by the two mechanisms: (i) high expression levels (*μ*_*p*_) are required for high noise amplitude 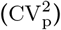 or (ii) lower expression levels coupled with lower noise.This trade-off leads to a ridge-like feature of nearly optimal models. The optimal fitness corresponds to a balance between increasing the mean protein number (*μ*_*p*_) and decreasing noise 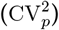. This optimal central dogma program strategy leads to significant overabundance.

**TABLE S1.** RLTO model parameters by figure. *Range* appears if a range of parameters is used. *NA* appears if the parameter value is irrelevant. *Local optimum* appears if the parameter values in optimized to maximize the growth rate.

| Figure number and panel: | Relative load: $\Lambda$<br>(No units) | Protein cost: $\lambda$<br>(molecules<sup>-1</sup>) | Protein threshold: $n_p$<br>(molecules) | Message number: $\mu_m$<br>(molecules/cell cycle) | Translation efficiency: $\varepsilon$<br>(No units) | Noise floor $C_0$<br>(No units) |
| --- | --- | --- | --- | --- | --- | --- |
| 2A | $10^{-5}$ | 10 | $10^2$ | Range | Range | 0 |
| 2C | Range | NA | Range | Local optimum | Local optimum | 0 |
| 4A | Weak dependence ($10^{-5}$) | NA | Range | Local optimum | Local optimum | 0 |
| 4B | Weak dependence ($10^{-5}$) | NA | Range | Local optimum | Local optimum | 0 |
| 4C | Weak dependence ($10^{-5}$) | NA | Range | Local optimum | Local optimum | 0 |
| 4D | Weak dependence ($10^{-5}$) | NA | Range | Local optimum | Local optimum | 0 |
| 5 | Weak dependence ($10^{-5}$) | NA | Range | Local optimum | Local optimum | 0 |
| S2A | $10^{-5}$ | 10 | $10^2$ | Range | Range | 0 |
| S2B | $10^{-5}$ | 10 | $10^2$ | Range | Range | 0 |
| S2C | $10^{-5}$ | 10 | $10^2$ | Range | Range | 0 |
| S2D | $10^{-5}$ | 10 | $10^2$ | Range | Range | 0 |
| S3A | Range | $10^2$ | Range | Local optimum | Local optimum | 0 |
| S3B | Range | $10^2$ | Range | Local optimum | Local optimum | 0 |
| S3C | Range | $10^2$ | Range | Local optimum | Local optimum | 0 |
| S3D | Range | $10^2$ | Range | Local optimum | Local optimum | 0 |
| S4A | Range | $10^2$ | Range | Local optimum | Local optimum | 0 |
| S4B | Range | $10^2$ | Range | Local optimum | Local optimum | 0 |
| S4C | Range | $10^2$ | Range | Local optimum | Local optimum | 0 |
| S4D | Range | $10^2$ | Range | Local optimum | Local optimum | 0 |
| S5 | $10^{-5}$ | NA | Range | Local optimum | 30 | 0.1 |
| S6A | Range | NA | Range | Local optimum | Local optimum | 0 |
| S7 | Weak dependence ($10^{-5}$) | NA | Range | Local optimum | Local optimum | 0 |
| S9A | Range | NA | Range | Local optimum | Local optimum | 0 |
| S12B | Weak dependence ($10^{-5}$) | NA | Range | Local optimum | Local optimum | 0 |

### 4. Methods: Central dogma optimization

#### a. Optimization of transcription and translation (eukaryotes)

The relative growth rate is:

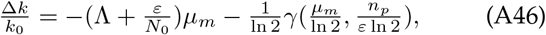

where *γ* is the regularized incomplete gamma function, which is the CDF of the gamma distribution and represents the probability of arrest due to gene *i*. (Note that this equation is identical to Eq. A45 but with the gene subscript *i* implicit.) We set the partial derivative of the growth rate with respect to message number equal to zero:

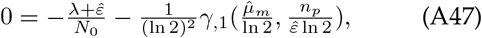

where we use the canonical comma notation to show which argument of *γ* has been differentiated. Next we differentiate with respect to the translation efficiency to generate a second optimization condition:

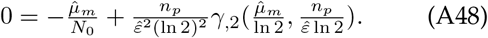

We will work in the large multiplicity limit where the overall metabolic load is much smaller than the metabolic load associated with any single gene: 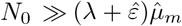. Next, we eliminate the threshold *n*_*p*_ in favor of the optimal overabundance:

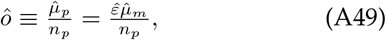

in both Eqs. A47 and A48. Eq. A48 can now be solved for the optimal translation efficiency:

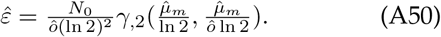

If we reinterpret *γ* as the CDF of the gamma distribution, we can rewrite this equation in terms of the gamma distribution PDF:

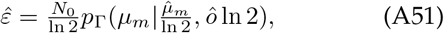

which will be the optimization equation for the translation efficiency.

To derive the optimization condition for the message number *μ*_*m*_, we substitute Eq. A50 into Eq. A47:

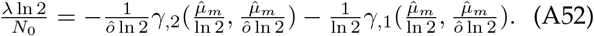

The two terms on the RHS can now be collected as the single partial derivative of message number *μ*_*m*_:

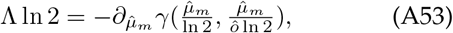

where the relative load is Λ ≡ *λ/N*_0_.

**FIG. S3.**
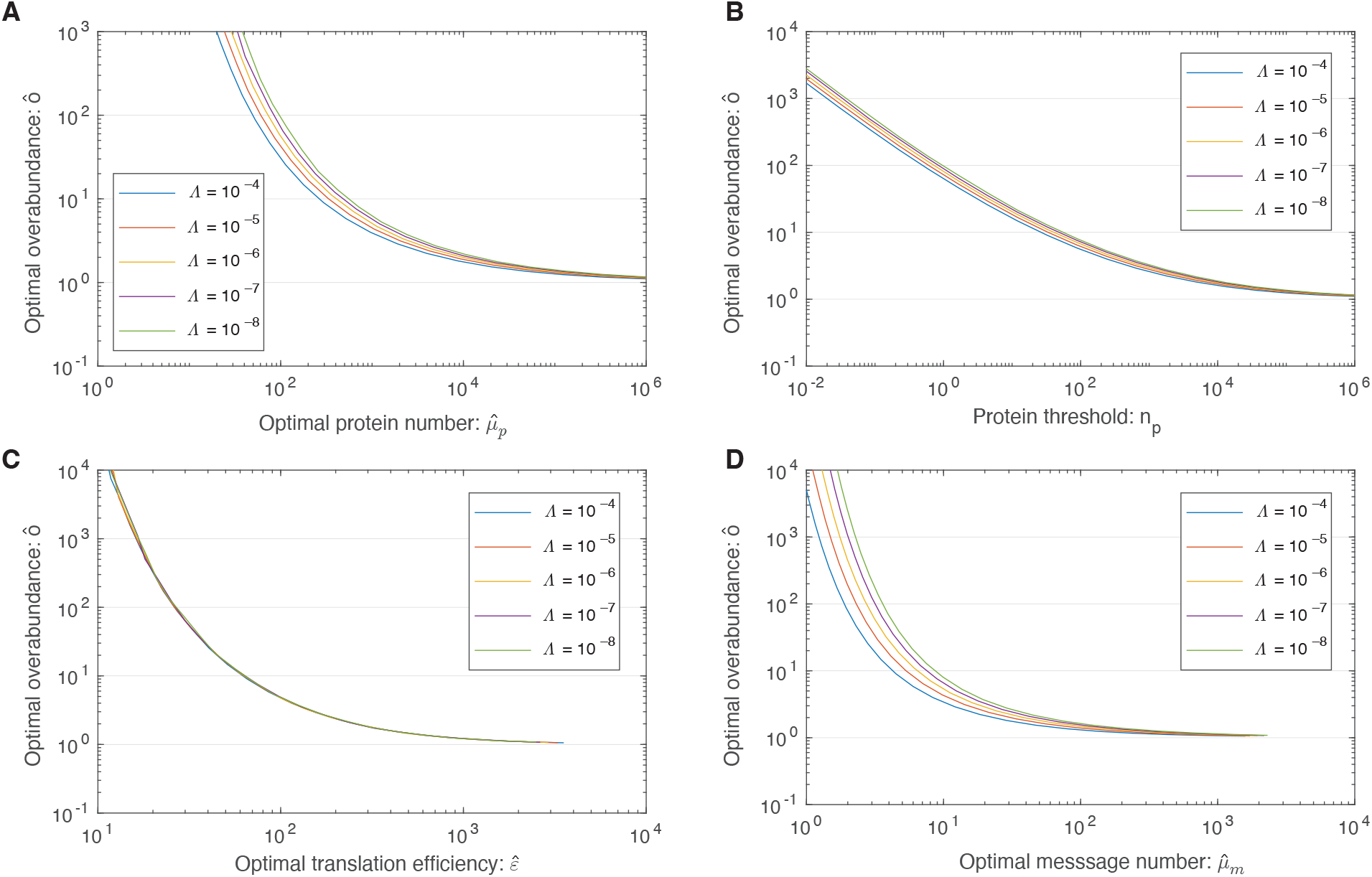
Four perspectives on the dependence of optimal overabundance on relative load Λ. All these calculations are performed for protein cost *λ* = 100 in order to give real numbers in molecules per cell. **Panel A: Overabundance as a function of protein number**. Overabundance decreases as protein number increases. These calculations are *λ dependent*. **Panel B: Overabundance as a function of the protein threshold**. Overabundance decreases as protein threshold increases. These calculations are *λ dependent*. **Panel C: Overabundance as a function of translation efficiency**. Overabundance decreases as translation efficiency increases. These calculations are *λ dependent*. **Panel D: Overabundance as a function of message number**. Overabundance decreases as message number increases. These calculations are *λ independent*.

The two optimization equations are summarized below:

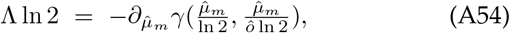

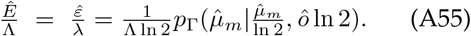

The optimal overabundance is shown for a range of relative loads in Fig. S3. The optimal translation efficiency and scaled translation efficiency are shown for a range of relative loads in Fig. S4.

#### b. Optimization of message number only

Consider the special case of optimizing the message number only at fixed translation efficiency. Eq. A47 is the condition; however, in this case it makes sense to adsorb both the message and protein metabolic load into a single metabolic load. The optimum message number satisfies the equation:

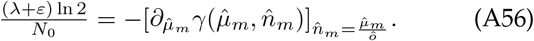

We define a modified relative load:

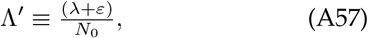

and substitute this into the optimum message number equation:

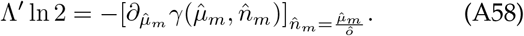

which is clearly closely related to Eq. A53.

We compare this modified expression to the original for optimum overabundance as a function of message number in Fig. 2C and demonstrate that the two make nearly identical predictions.

**FIG. S4.**
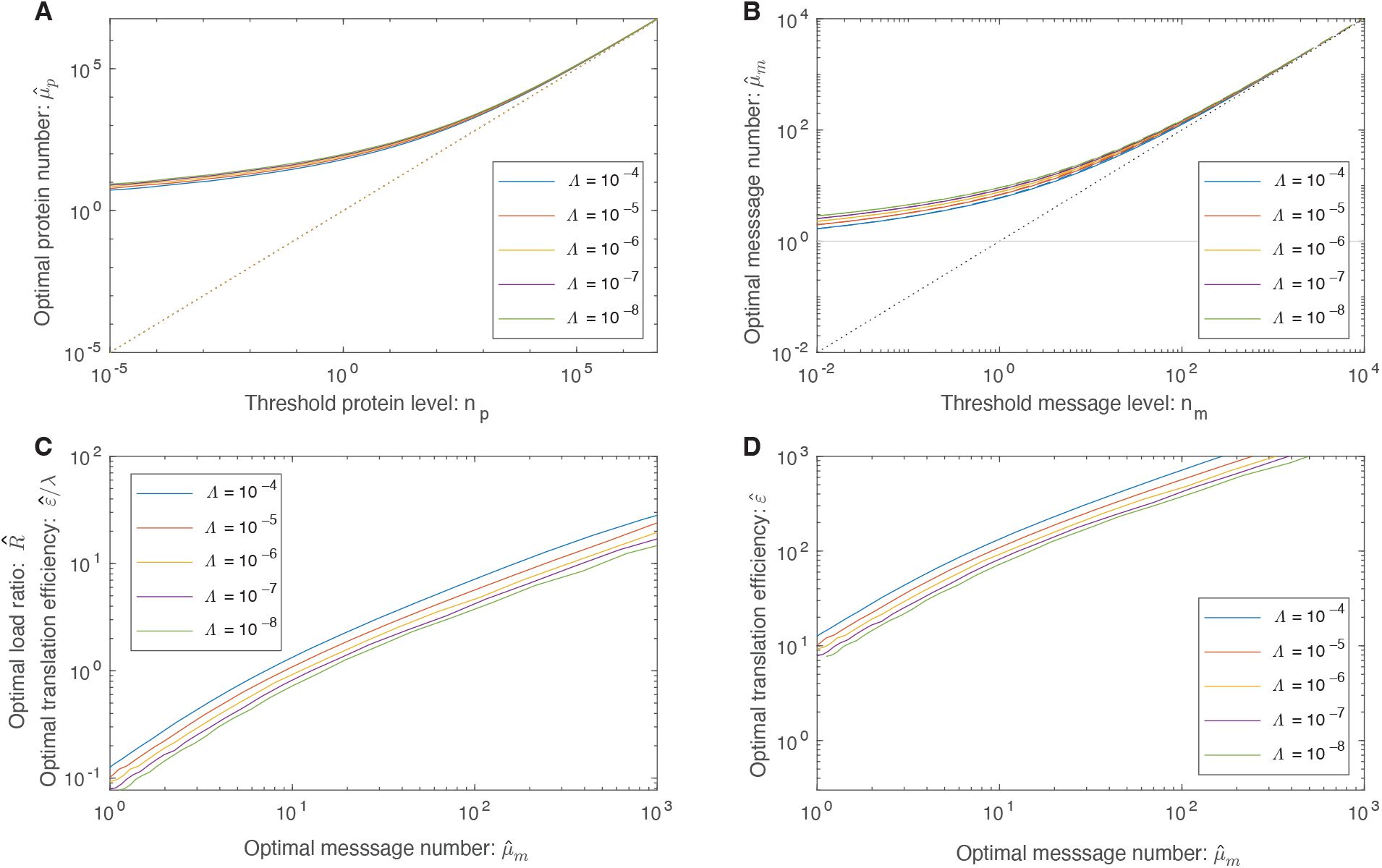
Four perspectives on load balancing. All these calculations are performed for protein cost *λ* = 100 in order to give real numbers in molecules per cell. **Panel A: Protein number versus protein threshold**. At high expression levels, the protein number tracks the protein threshold; however, the one message rule forces the protein number to threshold for low expression levels. These calculations are *λ dependent*. **Panel B: Message number versus message threshold**. At high expression levels, the message number tracks the message threshold; however, the one message rule forces the message number to a threshold close to *μ* _*m*_ = 1 for low expression levels. These calculations are *λ independent*. **Panel C & D: Message number versus translation efficiency**. The optimal translation efficiency grows almost linearly with the optimal message number. The scaled translation efficiency 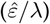 is independent of *λ* while the translation efficiency 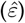 is dependent on *λ*. The ratio *ε/λ* has a second interpretation: the load ratio *R. R* is defined as the metabolic cost of translation over transcription of the gene.

#### c. Inclusion of the noise floor

In bacterial cells, the noise is dominated by the noise floor for high expression genes. Including the noise floor, the coefficient of variation squared is ([9] and Sec. D):

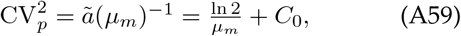

where *C*_0_ = 0.1 for bacterial cells [9]. In spite of the addition of noise from the noise floor, the observed distribution of protein number is still well described by the gamma distribution [9]; however, we need to modify the statistical parameters to account for the noise floor. (See the definition of the *statistical noise model* in Sec. A 1 b.) The modified gamma parameters are:

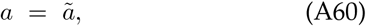

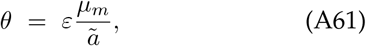

chosen such that the noise is determined by Eq. A59 but the protein number remains:

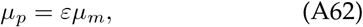

the product of the message number and translation efficiency.

The qualitative effect of the noise floor is to increase the noise, especially for low-copy messages. Above *μ*_*m*_ = 7 messages, the noise is dominated by the noise floor. Increases in transcription above this point have little effect on reducing the noise. As a consequence, the overabundance stays high, even for high copy messages. We compare this modified expression to the original for optimum overabundance as a function of message number in Fig. 2C and demonstrate that bacterial cells are predicted to have much higher overabundance at high expression levels.

**FIG. S5.**
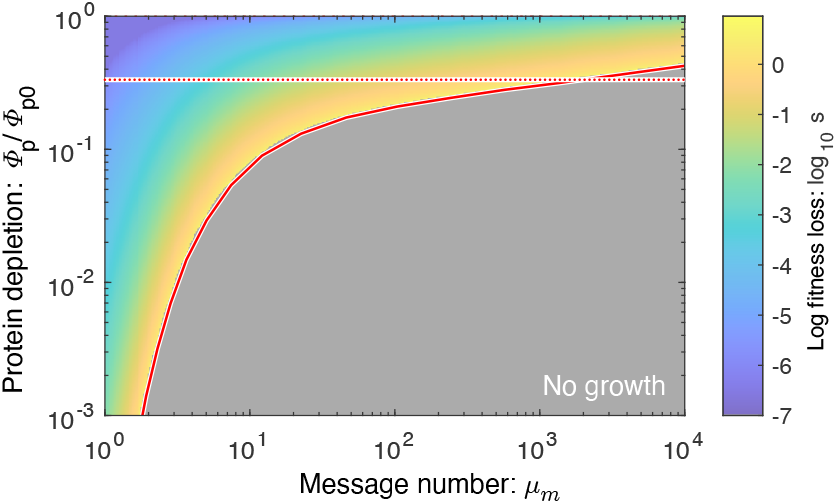
Optimal expression levels are buffered. The predicted fitness loss as a function of protein depletion level and message number for bacterial cells (including the noise floor). Due to the overabundance phenomenon, all proteins are buffered against depletion, but low-expression genes are particularly robust due to higher overabundance. The solid red line represents 1*/o*, and predicts the range of depletion values for which cell growth is predicted. The dotted red line represents a three-fold depletion.

### 5. Discussion: Understanding the rationale for overabundance

Essential protein overabundance is the signature prediction of the RLTO model. Its mathematical rationale is the highly-asymmetric fitness landscape. To understand why we expect this rationale to be generic, consider the form of the optimization condition for message number:

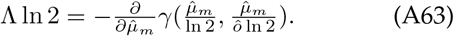

The growth rate is maximized when the probability of slow-growth (*e*.*g*. arrest) is roughly equal to the relative load of adding one more message. Since the cell makes roughly 10^5^ messages per cell cycle, the relative load is extremely small and therefore the probability of slow growth must be as well. Making this probability very small requires vast overabundance for the inherently-noisy, low-expression proteins.

The reason we expect the RLTO model protein abundance predictions to be robust is that we generically expect the fitness cost of overabundance to be small due to the high multiplicity (*i*.*e*. total number of genes); whereas, the fitness cost of arrest of essential processes is very high.

### 6. Discussion: RLTO predicts larger overabundance in bacteria

There are two distinctive features of bacterial cells that could affect the model predictions: (i) the translation efficiency is constant [22] and bacterial gene expression is subject to a large-magnitude noise floor that increases the noise for high-expression genes [9]. The optimization of message number at fixed translation efficiency does result in a slightly modified optimization condition for the message number (Sec. A 4 b); however, the predicted overabundance is only subtly perturbed (Fig. 2C). In contrast, the noise floor increases the predicted over-abundance, especially for high-expression proteins. As a result, the RLTO model predicts that the vast majority of bacterial proteins are expressed in significant over-abundance. (See Fig. 2C.)

### 7. Discussion: RLTO predicts proteins are buffered to depletion

A principle motivation for our analysis is the observation that many protein levels appear to be buffered. To explore the prediction of the RLTO model for protein depletion, we first computed the optimal message numbers and translation efficiencies for a range of protein thresholds. To model the effect of protein depletion, we computed the change in growth rate as function of protein depletion (equivalent to a reduction of the translation efficiency relative to the optimum.) The growth rate is shown in Fig. S5 for the RLTO model with parameters representative of a bacterial cell. (See Sec. A 4 c.)

In general, the RLTO model predicts that protein numbers have very significant robustness (*i*.*e*. buffering) to protein depletion. This is especially true for low expression proteins that are predicted to have the largest overabundance. For these genes, even a ten-fold depletion leads to very subtle reductions in the growth rate. For a three-fold reduction in the growth rate, only the very highest-expression genes (*e*.*g*. ribosomal genes) are expected to lead to qualitative phenotypes.

### 8. Results: Detailed development of load balancing

#### a. Prediction of the optimal load ratio

The two-stage amplification of the central dogma implies that the expression and noise levels can be controlled independently by the balance of transcription to translation. How does the cell achieve high and low gene expression optimally, and how does this strategy depend on the message cost?

To understand the optimization, we first define the load ratio *R* for a gene as the metabolic cost of translation relative to transcription:

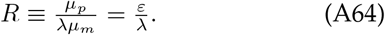

In Sec. A 4, we show that the optimal load ratio is:

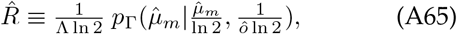

where *p*_Γ_ is the PDF of the gamma distribution. The optimal load ratio is shown in Fig. S4C.

The dependence of the optimal load ratio 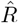 on Λ is extremely weak, but it is strongly dependent on message number. As a result, for low transcription genes (*μ*_*m*_ ≪ 10), the metabolic load is predicted to be dominated by transcription; whereas, for highly transcribed genes (*μ*_*m*_ ≫ 10), the metabolic load is dominated by translation. These predictions are robust since they are independent of the relative load Λ.

#### b. Measurements of the load ratio

Unfortunately, there is somewhat limited data to which to compare the model. The best source we found was Kafri *et al*. [43] who analyzed the differences in fitness between transcription and transcriptional-and-translation of a fluorescent protein driven by the pTDH3 promoter in yeast. This promoter is one of the strongest in yeast. Based on the RLTO model, we would predict this promoter to have a very high translation efficiency and therefore a large load ratio; however, the translation efficiency is much lower than one would predict based on a global analysis and likewise its load ratio is roughly unity, which based on the smaller than expected translation efficiency is broadly consistent with our expectations. A satisfactory test of this prediction will require larger-scale measurements that probe more representative genes.

### 9. Results: Translation efficiency is predicted to increase with transcription

Now that we have defined the optimal load ratio (Eq. A65), the equation for optimal translation efficiency can be written concisely:

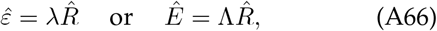

where 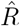 depends weakly on the relative load Λ. The RLTO model predicts that optimal partitioning of amplification between transcription (gain *μ*_*m*_) and translation (gain *ε*) has two important qualitative features: (i) As the message cost (*λ*) rises, the optimal translation efficiency increases in proportion. (ii) The optimal translation efficiency is also approximately proportional to message number 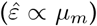. (See Fig. 4A.) Therefore, the RLTO model predicts that low expression levels should be achieved with low levels of transcription and translation, whereas high expression genes are achieved with high levels of both. We call this relation between optimal transcription and translation *load balancing*.

### 10. Results: RLTO predicts that message number responds to message cost

We will first focus on analyzing the implications of the message cost dependence in Eq. A66. At a fixed load ratio, Eq. A66 clearly implies that the translation efficiency increases as the message cost *λ* increases; however, the message number (and load ratio) also respond to compensate to changes in *λ*. To probe the dependence on message cost in an experimentally relevant context, consider optimal message numbers in a reference condition (relative load Λ_0_) relative to a second perturbed condition (relative load Λ). The predicted relation between the optimal messages numbers is shown in Fig. S6A. The resulting relation between the optimal message numbers is roughly linear on a log-log plot, predicting the approximate power-law relation:

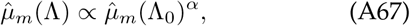

describing a non-trivial global change in the regulatory program.

### 11. Results: RLTO predicts the yeast global regulatory response

To test the RLTO predictions, we compared the relative message numbers for yeast growing under phosphate depletion, which increases the message cost [43], to a reference condition [44]. As predicted, the relative transcriptome data was well described by a power law (Eq. A67) and the observed slope was smaller than one: 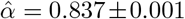, as predicted by the increased message cost. See Fig. S6B.

The observation of this large-scale regulatory change has an important implication: This response supports a nontrivial hypothesis that the RLTO model not only can predict how the cell is optimized in an evolutionary sense, but can predict global regulatory responses as well.

Note that Metzl-Raz *et al*. [44] also explored conditions that increased the cost of protein; however, here the predictions of the model are ambiguous. The complication arises due to the observed decrease in size of the cells in the experimental condition, which decreases the total metabolic load *N*_0_. As discussed above, the relative load Λ = *λ/N*_0_ is the key determinant in Eq. A67; however, even as the relative cost of transcription *λ* decreases in the experimental condition, the total metabolic load *N*_0_ also decreases, making no clear prediction about how the relative load Λ changes.

### 12. Methods: Prediction of the proteome fraction

#### a. Parameter-free prediction of proteome fraction

**FIG. S6.**
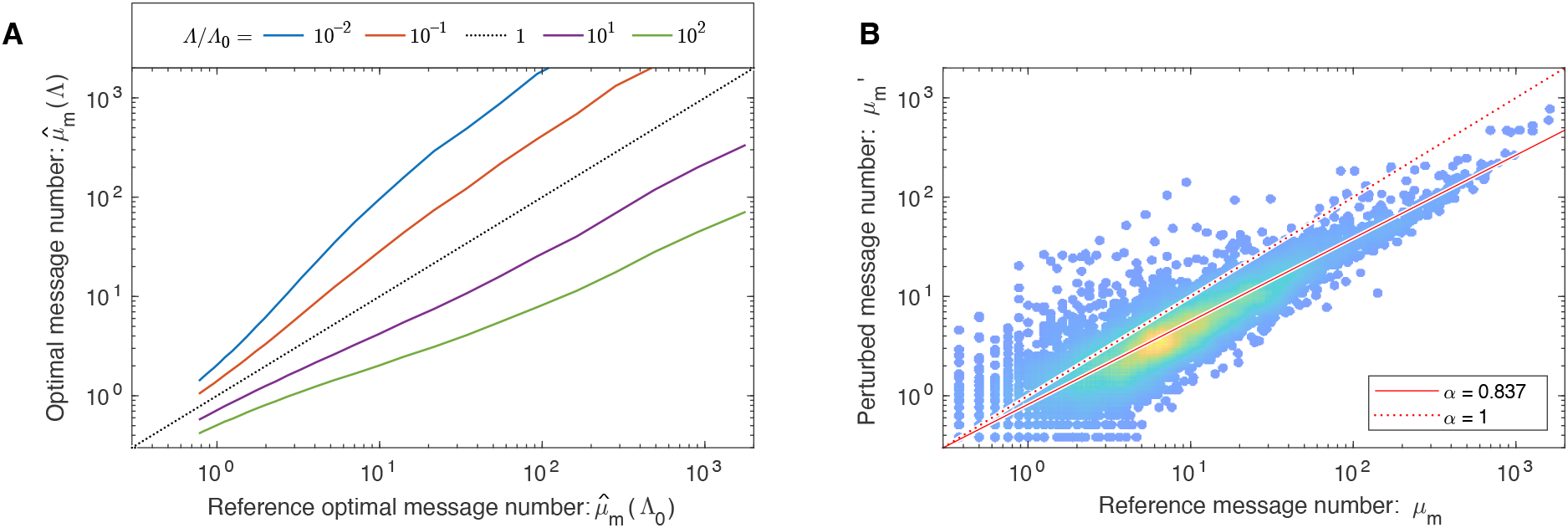
The message cost affects transciption genome wide. **Panel A: Message number decreases with increased relative load Λ.** The optimal message number responds to changes in the message cost. The RLTO model predicts an approximate power-law relation (linear on a log-log plot) between message numbers. **Panel B: A power-law relation is observed**. To test whether central dogma regulation would adapt dynamically as predicted, we analyzed the relation between the yeast transcriptome under reference conditions and phosphate depletion (perturbed), which increases the message cost [43]. (Data from Ref. [44].) As predicted by the RLTO model, a global change in regulation is observed, which generates a power-law relation with scaling exponent *α* = 0.837 *±* 0.01. The observed exponent is smaller than one, as predicted by an increased relative load Λ.

We now turn our focus to an analysis of the implications of *load balancing*: the message number dependence of the optimal translation efficiency (Eq. A66). The most direct test of this prediction is measuring the relation between proteome fraction and message number. The RLTO model predicts proteome fraction (Eq. A34):

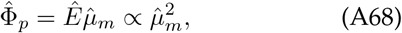

where *μ*_*m*_ is the observed message number and the optimal relative translation efficiency is predicted by Eq. A66. The proportionality is only approximate but gives important intuition for how protein number depends on message number in the RLTO model, in contrast to a constant-translation-efficiency model: Φ_*p*_ ∝ *μ*_*m*_. To compare these predictions to protein abundance measurements, we will renormalize the protein fraction to be defined relative to total protein number rather than *N*_0_. This renormalization eliminates the Λ dependence to result in a parameter-free prediction of the proteome fraction.

#### b. RLTO: proteome fraction

Starting from Eq. A96, clearly:

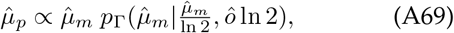

which can be used to predict the proteome fraction (where we have restored the explicit gene *i* subscript):

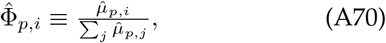

where the second subscript is the gene index. To predict the proteome fraction, we computed the proportionality constant *C*:

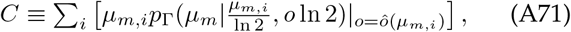

where the message numbers *μ*_*mi*_ for gene *i* are the experimentally observed message numbers, the implicit *o*_*i*_ values are predicted by the RLTO model (Eq. A53) for message number *μ*_*mi*_ and the sum index *i* runs over all genes. The predicted optimal proteome fraction is:

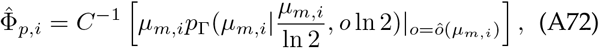

which generates the predicted solid curves shown in Fig. 4BCD.

#### c. Constant-translation-efficiency model: proteome fraction

For the constant translation efficiency model, we define the normalization:

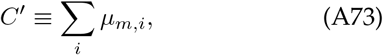

and the predicted proteome fraction is:

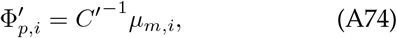

which generates the predicted dotted curves shown in Fig. 4BCD.

#### d. Sources of experimental data for proteome fraction analysis

For *E. coli* data, the protein abundance data was generated by mass spec measurements and the message abundance data was from [22]. For the yeast data, the protein abundance data is measured by mass spec and message abundances are determined by [21]. For the mammalian data, we used mouse data. The protein abundance data is measured by mass-spec and message abundances are determined by [23].

We estimated the message number *μ*_*m*_ as described in Sec. C 2 c. For the mouse data, the study provided message lifetimes, the cell cycle duration and abundances in molecules per cell [23]. For the *E. coli* and yeast, the total number of proteins, messages *etc*, cell cycle duration and message lifetimes for each organism and their sources are described in Tab. S2.

### 13. Results: Load balancing is observed in eukaryotic cells

A non-trivial prediction of the RLTO Model is that translation efficiency and message number should be roughly proportional. Qualitatively, this strategy allows expression levels to be increased while distributing the added metabolic load between transcription, which reduces noise, and translation, which does not affect the noise. We predict the optimal translation efficiency versus message number which matches the observations in eukaryotic cells (Fig. 4BC). However, in *E. coli*, the translation efficiency and message number are *not* strongly correlated (Fig. 4D). Why does this organism appear not to load balance? We demonstrate that the observed translation efficiency is consistent with the RLTO model, augmented by a ribosome-per-message limit. (See Sec. A 15.) Hausser *et al*. have proposed just such a limit, based on the ribosome footprint of mRNA molecules [10]. (See Sec. A 17.) Although this augmented model is consistent with central dogma regulation in *E. coli*, it is not a complete rationale. This proposed translation-rate limit could be circumvented by increasing the lifetime of *E. coli* messages which would increase the translation efficiency. Why the message lifetime is as short as observed will require a more detailed *E. coli*-specific analysis.

### 14. Discussion: Relation between load balancing and previous results

Hausser *et al*. have previously performed a more limited analysis of the trade-off between metabolic load and gene-expression noise [10]. In this section, we will provide some more context into the differences between the two approaches.

Hausser *et al*. assume a symmetric (not an asymmetric) fitness landscape and consider only the metabolic cost of transcription (but not translation). Their model depends on two (not one) gene-specific parameters: an optimal protein number and a sensitivity, which defines the curvature of the fitness [10]. The authors maximize fitness with respect to the transcription rate (but not the translation rate) and the condition they derive depends on the two (not one) unknown, gene-specific parameters. As a result, this condition is not predictive of global regulatory trends without non-trivial, genespecific measurements or assumptions about the unknown sensitivity.

The Hausser model assumes that the growth rate has the form:

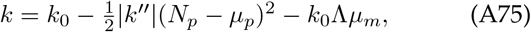

where *k* is the growth rate, and we have rewritten the form of the fitness to better match our own definitions. Here *k*^*′′*^ is the second derivative of the growth rate at the optimal protein number *μ*_*p*_ and *N*_*p*_ is the stochastic protein number. If we take the expectation with respect to the protein number, we get:

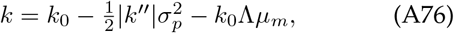

and substituting the noise model for the variance of the protein number gives:

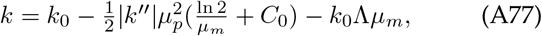

where *C*_0_ is the noise floor, and we have assumed the mean protein number is optimal (*μ*_*p*_). If we maximize the growth rate with respect to *μ*_*m*_, we get the following condition on the optimal message number:

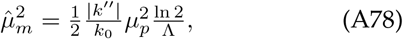

which depends on the unknown curvature *k*^*″*^. To make global predictions about how transcription and translation are related, some added assumptions are necessary to describe how *k*^*″*^ scales with protein abundance.

To illustrate how this expression does not make explicit global predictions, let’s consider a number of plausible possibilities. First, we will assume that *k*^*″*^ is independent of *μ*_*p*_ and on average all proteins are equally sensitive to changes in protein number. In this case, we find:

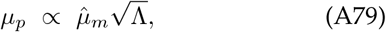

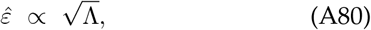

implying a constant translation efficiency which is inversely proportional to the square root of the relative load.

Alternatively, we can assume that 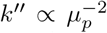 and, on average, the cell is equally sensitive to changes in the relative number of proteins (*i*.*e*. Δ*p/μ*_*p*_), regardless of expression level. In this case,

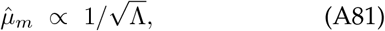

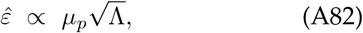

implying a constant message number, irrespective of expression level, and a translation efficiency that is proportional to expression level.

Finally, we will assume that 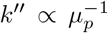, which is the intermediate case. Here:

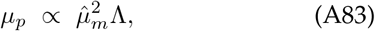

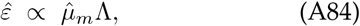

implying that translation efficiency should increase with message number, analogous to our prediction. It appears that Hausser *et al*. implicitly also favor this model, since they define their sensitivity parameter to include a power of protein number *μ*_*p*_. They justify this assumption by arguing that since 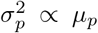, it makes sense to define the *sensitivity to noise* to include a factor of *μ*_*p*_ [10]. At best, this is somewhat fuzzy logic since, as we demonstrate in the paper, Eq. A84 implies that the protein variance does not scale 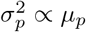!

**FIG. S7.**
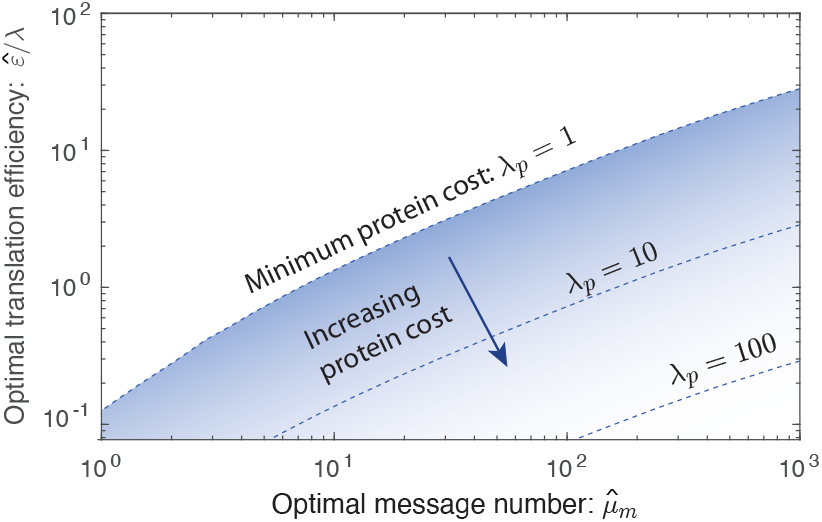
Increased protein cost decreases optimal translation efficiency. A protein cost of *λ* _*p*_ = 1 corresponds to the metabolic cost of protein synthesis only, and is the minimum protein cost. For larger protein costs, the optimal translation efficiency is lower. As a result, the *λ* _*p*_ = 1 curve represents an upper bound of the optimal translation efficiency.

The authors also propose a lower limit on the translation-transcription ratio; however, their limit is dependent on the noise floor, which only affects genes with the highest transcription rates in eukaryotic cells. The implementation of a more appropriate estimate of the noise, relevant for the vast majority of genes, does not lead to the same limit.

### 15. Methods: Analysis of translational limits & gene-specific load analysis

To explore the consequences of a protein-specific load, we can modify the metabolic load term in the growth rate equation (Eq. A46):

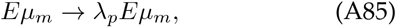

which includes an additional parameter: the protein cost *λ*_*p*_, which is 1 if the fitness cost is equal to the metabolic load and greater than one if the cost is higher. We will also treat the metabolic load per message *λ* as a gene-specific parameter in this section only. The optimization can be repeated for this augmented model.

To analyze the effect of increased protein load, we modify Eq. A46:

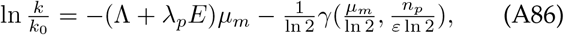

to contain the supplemental load factor *λ*_*p*_ which is unity if the only protein load is metabolic and *λ*_*p*_ *>* 1 if there is additional load (*e*.*g*. toxicity). The optimization conditions (Eqs. A47 and A48) become:

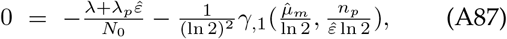

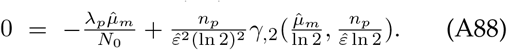

Using the same algebraic approach as before, we can derive the same optimal overabundance and load equations:

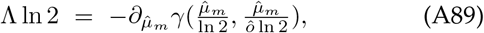

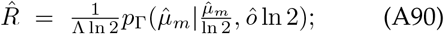

however, the relation between the load and the translation efficiency now has an extra factor: *λ*_*p*_:

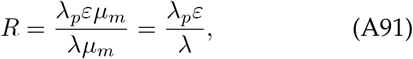

representing the modified total load ratio.

### 16. Results: Increased protein-specific cost reduces the optimal translation efficiency

The relation between the overabundance and message number is unchanged (Eq. A63). This result can be rationalized in the following way: The optimal over-abundance is determined by the noise which is determined by message number only. This relation is unaffected by the added parameter *λ*_*p*_. However, the optimal translation efficiency is affected:

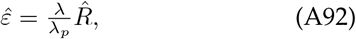

where 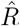 is the optimal load ratio, defined by Eq. A66. The optimal curves are shown in Fig. S7.

How do these added considerations affect the RLTO predictions? First, we consider message and protein length. What are the optimal translation efficiencies for two proteins, one ten times the length of the other, at fixed protein number? In this case, we will assume that both the transcriptional cost (*λ*) as well as the translational cost (*λ*_*p*_) increase tenfold. These increases cancel, resulting in the same optimal translation efficiency since it is only the relative cost of transcription to translation that is determinative of the translation efficiency.

Now consider a tenfold protein-specific increase in protein cost at fixed message cost and fixed protein number. The message number and translation efficiency would change by compensatory factors of 10:

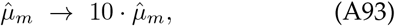

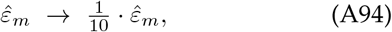

to maintain the protein number.

Returning to our original motivation, we can understand how genes with a higher protein-to-message cost migrate downwards and rightwards off the optimal *λ*_*p*_ = 1 curve, predicting a cloud versus a narrow strip in proteome fraction measurements shown in Fig. 4. If the relative load Λ were directly measured, we would expect the predicted optimal translation efficiency curve for *λ*_*p*_ = 1 to lie at the top edge of the observed data cloud rather than the bisecting it. This bisection is the consequence of fitting an effective relative load parameter to the abundance data in the unaugmented RLTO model.

### 17. Discussion: Translation limits in *E. coli*

A critical assumption in the RLTO model to this point has been that the optimal central dogma parameters are realizable in the cell; however, translation can be limited by a number of different mechanisms. The superior performance of the constant-over the optimal-translation-efficiency model in *E. coli* (Fig. 4D) suggests that this assumption may not be satisfied for bacteria. How do translation limits affect the model phenomenology?

When considering possible limits on translation, there are two natural mechanisms: (i) ribosome-number limit, where the number of ribosomes in the cell limits translation and (ii) a ribosome-per-message limit, where the number of ribosomes per message is limiting. Assuming the ribosome-number-limit mechanism, the original unconstrained optimization problem can be recast as a constrained optimization problem where the protein cost *λ*_*p*_ is reinterpreted as a Lagrange multiplier to constrain the number of proteins translated (*e*.*g*. [45]). In spite of this reformulation, we would still predict the same functional form for the coupling between the optimal translation efficiency and message number. (*I*.*e*. it is still optimal to have a higher translation efficiency for highly-expressed genes even if the total number of proteins is fixed.) Therefore, the ribosome-number-limit mechanism cannot be the rationale for the constant translation efficiency observed in *E. coli*.

Assuming the ribosome-per-message-limit mechanism, we limit the translation efficiency to a restricted range of values. If the unconstrained optimum lies above this range, the optimum is at the maximum limiting value. If the unconstrained optima for all genes lie above the realizable range, the model predicts a translation efficiency uncoupled from the message number, as observed. These predictions are consistent with the observed central dogma regulatory program in *E. coli*. In added support of this hypothesis, Hausser *et al*. have argued that *E. coli* translates close to just such a ribosomeper-message limit as a consequence of the finite ribosome complex footprint on a message [10].

### 18. Methods: Estimate of the message cost and metabolic load

We can estimate the message cost *λ* from the known total protein number for yeast and mammalian cells. (For *E. coli* this estimate is not possible since the protein cost in not determinative of the translation efficiency.)

The optimal translation efficiency for gene *i* is (Eq. A51):

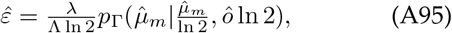

and therefore the optimal protein number for gene *i* is:

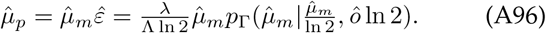

We define the normalization constant *A* (where we restore the explicit gene *i* subscript):

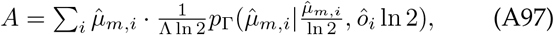

where we have restored the explicit gene *i* index running over all genes. Now, by summing Eq. A96, over all genes, we derive an expression for the total protein number 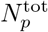 in terms of the message cost *λ* and the normalization constant *A*:

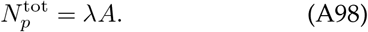

Solving for the protein cost results in the estimate:

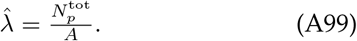

This message cost estimate 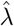 can then be plugged into the metabolic load definition:

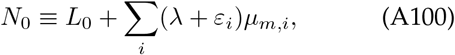

to estimate its size:

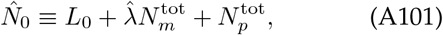

where we have ignored the non-protein and non-message contributions to the load (*L*_0_ = 0).

#### a. Detailed protocol

We first estimate the message numbers, as described in Sec. C 2 c, from data. For each gene *i*, we set the optimal message number equal to the observed message number and then compute the optimal overabundance from the message number using Eq. A54. (Since the result is independent of the assumed Λ value, we set an arbitrary initial value of Λ = 10^*−*5^.) We then use these single gene optimal message number and overabundances to compute *A* using Eq. A97. In Eqs. A99 and A101, we use the 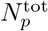 from Tab. S2. 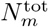 is computed by summing the estimated message numbers.

#### b. Estimate the message cost and metabolic load in yeast

In yeast, the estimates are:

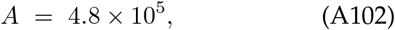

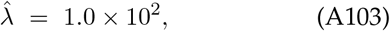

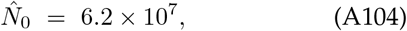

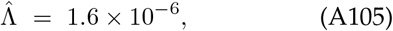

where the data sources are described in detail in Sec. C 1 b.

#### c. Estimate the message cost and metabolic load in human cells

In human cells, the estimates are:

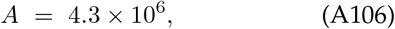

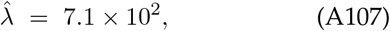

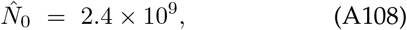

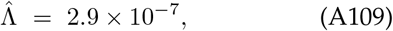

where the data sources are described in detail in Sec. C 1 c.

#### d. Estimate of the modified relative load in bacterial cells

In bacterial cells, we will assume a constant translation efficiency model. We therefore use the modified relative load formula (Eq. A57) to estimate Λ^*′*^. We will assume that the load is dominated by proteins and messages:

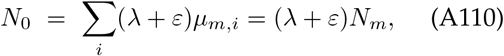

where *N*_*m*_ is the total number of messages. We can then solve this equation for Λ^*′*^:

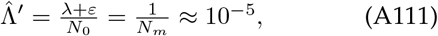

based on the total message number estimate for *E. coli*. (See Tab. S2.)

## Appendix B: Model robustness & exploring alternatives to RLTO

In this section, we investigate the phenomenology of three different single-cell growth rate functions to determine what model features result in overabundance. We consider an *arrest model* (the RLTO model), a *slow-growth model*, and a *symmetric model*.

**FIG. S8.**
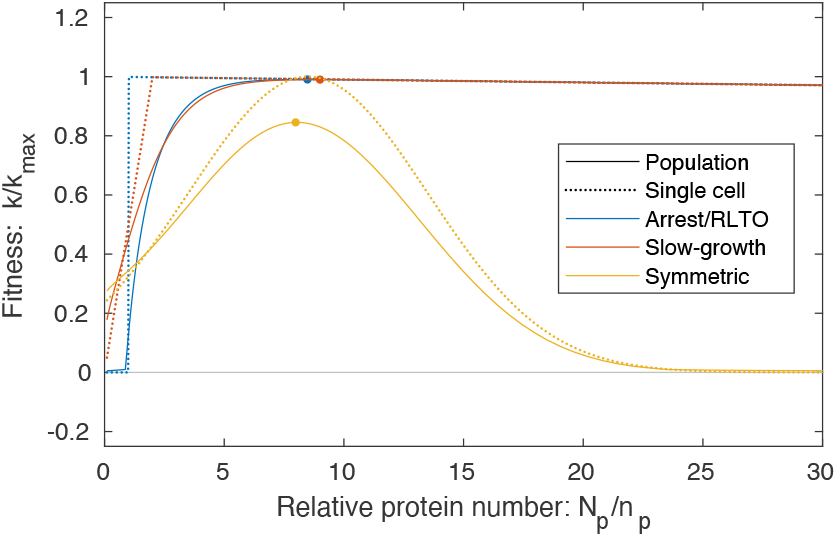
Exploring the mathematical mechanism of over-abundance. Single-cell and population growth rate are compared for three different models: arrest (RLTO), slow-growth, and symmetric models. In the arrest model (RLTO), the growth rate goes to zero below threshold protein level *n*_*p*_. In the slow-growth model, the growth rate transitions continuously to zero as the *N*_*p*_ is depleted below *n*_*p*_. In both the arrest and slow-growth models, there is a small negative slope above the threshold corresponding to the metabolic load. In the symmetric model, the fitness cost is symmetric about the optimum. Both the threshold-like and slow-growth models are optimized at mean expression levels *μ* _*p*_ far exceeding the threshold level *n*_*p*_. This is a consequence of the highly-asymmetric dependence of the fitness on protein number *N*_*p*_. This leads to the phenomenon of protein overabundance. In contrast, the symmetric model is optimized in close proximity to its single-cell optimum.

### 1. Methods: Defining alternative models

In each case, we will assume that the protein number is described by a gamma distribution:

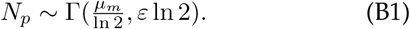

We will assume the cell-cycle duration *T* is determined by this stochastic protein number *N*_*p*_ and then compute the population growth rate using Eq. A36 for a range of different message numbers *μ*_*m*_. In each case, *τ*_0_ = 1*/N*_0_, *N*_0_ = 10^5^, *ε* = 30, *n*_*p*_ = *ε* ln 2. The mean expression level is *μ*_*p*_ = *μ*_*m*_*ε*.

#### a. Model 1: Arrest (RLTO) model

The arrest (RLTO) model has cell cycle duration:

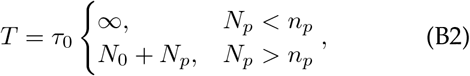

where protein expression below threshold *n*_*p*_ results in growth arrest.

#### b. Model 2: Slow-Growth model

In the slow-growth model, we imagine two processes: (i) checkpoint process X and (ii) other processes. The cell will divide after whichever process finishes last. Other processes will finish after time predicted by the metabolic load, identical to the threshold model defined above. However, we model checkpoint process X as the completion of a fixed amount of activity in an irreversible process. We will therefore assume it will take a time inversely proportional to the amount of enzyme X (*N*_*p*_). The amount of activity is set by effective threshold *n*_*p*_:

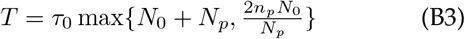

such that *n*_*p*_ defines the level of protein required to make the growth rate half the metabolic limit.

Unlike the arrest model, cell growth slows but does not stop for *N*_*p*_ *< n*_*p*_. This model will test whether the results of the RLTO model are an artifact of the assumed arrest-based slow growth.

#### c. Model 3: Symmetric model

For the symmmetric model, we choose the model parameters such that the single-cell optimum was close to the other models: *n*_0_ = 8.5, *σ*_*n*_ = 5. The cell-cycle duration is:

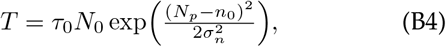

such that the noise-free growth rate will be Gaussian is *N*_*p*_.

### 2. Results: Overabundance is a robust prediction

The growth rates as a function of the mean expression level *μ*_*p*_ are shown in Fig. S8. The symmetric model has a population optimum in close proximity to its single-cell optimum, as we intuitively expect. However, both the arrest (RLTO) model and the slow-growth model have optima far above the threshold number *n*_*p*_. We therefore conclude that it is fitness asymmetry rather than growth arrest that is responsible for the overabundance phenomenon.

Why doesn’t growth arrest of a sub-population lead to a stronger effect than the same sub-population growing slowly? In Ref. [29], we showed that the population doubling time 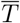 can be understood as the exponential mean of the stochastic cell-cycle duration:

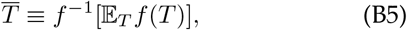

where E_*T*_ is the expectation over the stochastic duration *T* and *f* (*t*) ≡ exp(*−kt*), where 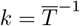 ln 2 is the population growth rate. Due to the functional form of *f* (*t*), any long cell cycles are exponentially suppressed in their contribution to the exponential mean. Therefore, low-probability extremely-long-duration cell cycles only contribute to the growth rate by reducing the fraction of growing cells.

## Appendix C: Quantitation of central dogma parameters for one-message-rule

The RLTO model predicts the *one-message-rule* for the lower threshold on transcription for essential genes. In this section, we use transcriptome data from the literature to test this prediction. We first describe the sources of the data (Sec. C 1), how the estimates are computed (Sec. C 2), the results (Sec. C 3) and discussion (Sec. C 4).

### 1. Methods: Selection of central dogma parameter estimates

The estimates for central dogma model parameters come from two types of data: (i) quantitative measurement of cellular-scale parameters for each organism (total number of messages in the cell, cell cycle duration, *etc*) and (ii) genome-wide studies quantitative of mRNA and protein abundance.

For the cellular-scale central dogma parameters, we relied heavily on an online compilation of biological numbers: BioNumbers [54]. This resource provides a collection of curated quantitative estimates for biological numbers, as well as their original source. In the interest of conciseness, we have cited only the original source in the Tab. S2, although we are extremely grateful and supportive of the creators of the BioNumbers website for helping us very efficiently identify consensus estimates for the parameters of the central dogma parameters.

For the selection of genome-wide studies on abundance, we used many of the same resources cited in BioNumbers as well as studies selected by a previous study of a quantitative analysis of the central dogma: Hausser *et al*. [10].

#### a. E. coli data

##### Message lifetimes

The message lifetimes (and median lifetime) were taken from a recent transcriptome-wide study by Chen *et al*. [36]. These investigators measured the lifetime in both rapid (LB) and slow growth (M9).

##### Noise

Taniguchi *et al*. have performed a beautiful simultaneous study of the proteome and transcriptome with single-molecule sensitivity [9]. Although we use the noise analysis data from this study for our supplemental analysis of *E. coli* noise, it is not the source for our *E. coli* transcriptome data due to the extremely slow growth of the cells in this study (150 minute doubling time), which is not consistent with the growth conditions for the other sources of data.

**TABLE S2.** Central dogma parameters for three model organisms with detailed references. Columns three through seven hold representative values for measured central-dogma parameters for the model organisms described in the paper. Each value is followed by a reference for its source.

| Model organism | Growth condition | Doubling time: $T$ | Message lifetime: $\tau_m = \gamma_m^{-1}$ | Message recycling ratio: $m = T/\tau_m$ | Total number of | | | Average | |
| --- | --- | --- | --- | --- | --- | --- | --- | --- | --- |
| | | | | | messages /cell: $N_{m/c}^{\text{tot}}$ | messages /cell-cycle: $N_m^{\text{tot}}$ | proteins: $N_p^{\text{tot}}$ | translation efficiency: $\varepsilon$ | translation rate: $\beta_p$ (h<sup>-1</sup>) |
| *Escherichia coli* (*E. coli*) | LB | 30 min [35] | 2.5 min [36] | 12 | $7.8 \times 10^3$ [46] | $9.4 \times 10^4$ | $3 \times 10^6$ [47] | 22 | 530 |
| | M9 | 90 min [35] | 2.5 min [36] | 36 | $2.4 \times 10^3$ [46] | $8.6 \times 10^4$ | $3 \times 10^6$ [47] | 24 | 580 |
| *Saccharomyces cerevisiae* (Yeast-haploid) | YEPD | 90 min [48] | 22 min [49] | 4 | $2.9 \times 10^4$ [50] | $1 \times 10^5$ | $5 \times 10^7$ [51] | $4 \times 10^2$ | 410 |
| *Mus musculus* (Mammalian-mouse) | Tissue | 27.5 h [23] | 15 h [23] | 1.8 | $1.7 \times 10^5$ [23] | $3 \times 10^5$ [23] | $3 \times 10^9$ [23] | $1 \times 10^4$ | 660 |
| *Homo sapiens* (Human) | Tissue | 24 h [52] | 14 h [53] | 1.7 | $3.6 \times 10^5$ [48] | $5 \times 10^5$ | $2 \times 10^9$ [47] | $4 \times 10^3$ | 120 |

##### mRNA abundance

Instead, we used data from the more recent Bartholomaus *et al*. study [46], which characterizes the transcriptome in both rapid (LB) and slow growth (M9).

##### Total cellular message number

This study was chosen since it was the source of the BioNumbers estimates of cellular message number in *E. coli* (BNID 112795 [54]).

##### Doubling time

The source of the doubling times for rapid (LB) and slow (M9) growth of *E. coli* comes from Bernstein [35].

##### Essential gene classification

The classification of essential genes in *E. coli* comes from the construction of the Keio knockout collection from Baba *et al*. [55].

#### Protein number

The total protein number in *E. coli* came from Milo’s recent review of this subject [47].

#### b. Yeast data

##### Message lifetimes

The message lifetimes (and median lifetime) were taken from Chia *et al*. [49].

##### Noise

The noise data was taken from the Newman *et al*. study, which used flow cytometry of a library of fluorescent fusions to characterize protein abundance with single-cell resolution [8].

##### mRNA abundance

The transcriptome data comes from the very recent Blevins *et al*. study [56].

##### Total cellular message number

There are a wide-range of estimates for the total cellular message number in yeast: 1.5 *×* 10^4^ [57] (BNID 104312 [54]), 1.2 *×* 10^4^ [58] (BNID 102988 [54]), 6.0 *×* 10^4^ [59] (BNID 103023 [54]), 2.6 *×* 10^4^ [50] (BNID 106763 [54]) and 3.0 *×* 10^4^ [60]. We used the compromise value of 2. *×* 10^4^.

##### Doubling time

The doubling time was taken from [48].

##### Protein number

The total protein number in yeast comes from Futcher *et al*. [51].

##### Essential gene classification

The classification of essential genes in yeast comes from van Leeuwen *et al*. [19].

##### Proteome abundance data

The proteome abundance data came from two sources: flow cytometry of fluorescent fusions from Newman *et al*. [8] as well as mass-spec data from de Godoy *et al*. [61].

#### c. Human data

##### Message lifetimes

The message lifetimes (and median lifetime) were taken from Yang *et al*. [53] who reported a median half life of 10 h which corresponds to a lifetime of 14 h.

##### mRNA abundance

The transcriptome data comes from the data compiled by the Human Protein Atlas [62], which we averaged over tissue types.

##### Total cellular message number

The total cellular message number in human comes from Velculescu *et al*. [63] (BNID 104330 [54]).

##### Doubling time

The doubling time was taken from [52].

##### Protein number

The total protein number in human came from Milo’s recent review of this subject [47].

##### Essential gene classification

The classification of essential genes in human comes from Wang *et al*. [20].

**FIG. S9.**
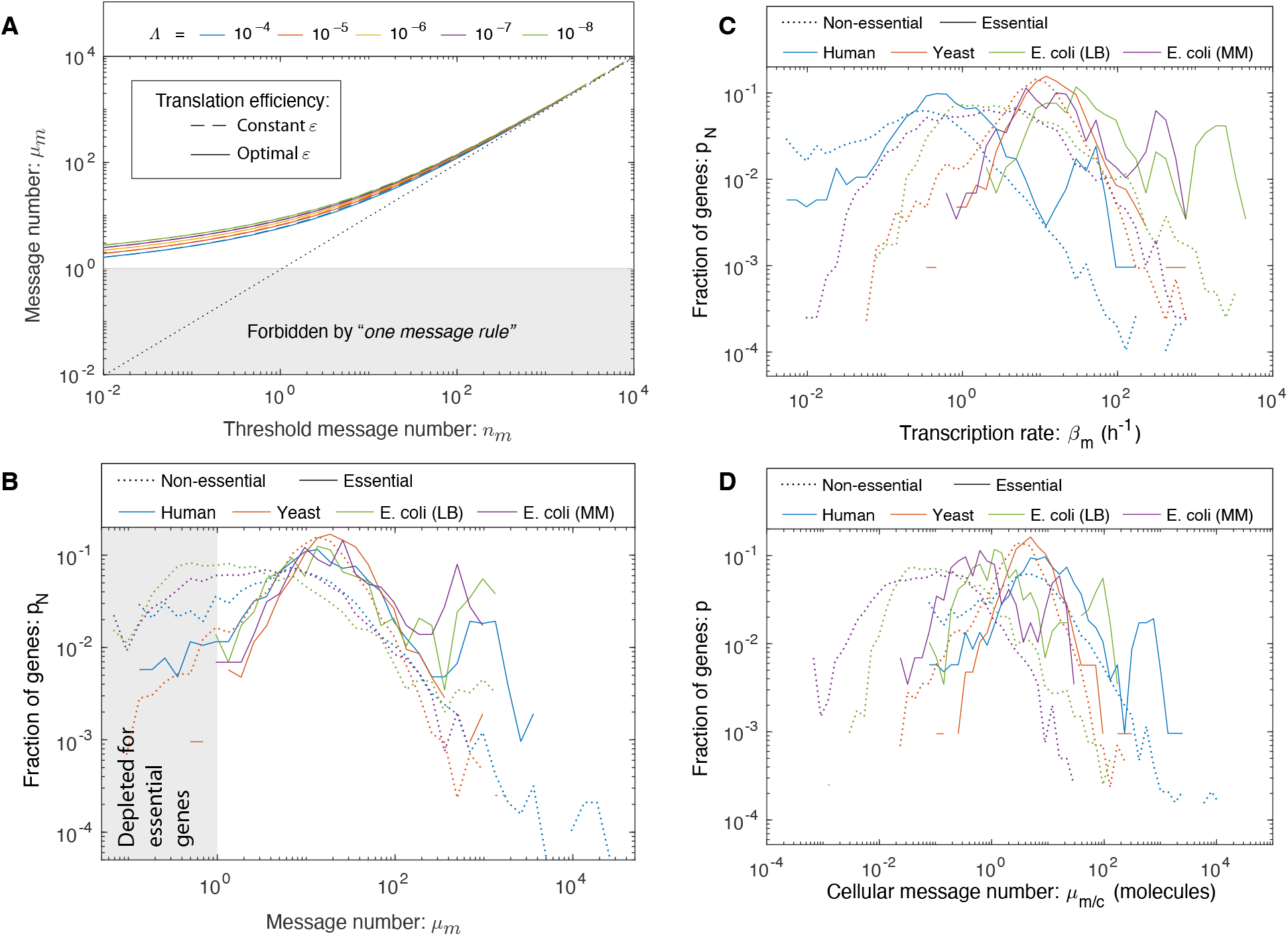
The one message rule. **Panel A: One-message-rule for essential genes.** For highly transcription genes (high *μ* _*m*_), little compensation for noise is required and the optimal message number tracks with the threshold message number *n*_*m*_. However, as the threshold message number approaches one (*n*_*m*_ → 1), the noise is comparable to the mean, and the optimal message number *μ* _*m*_ increases to compensate for the noise. As a result, a lower threshold of roughly one message per cell-cycle is required for essential genes. This threshold is predicted for both fixed (dashed) and optimized translation efficiency (solid). The threshold is weakly dependent on relative load Λ. **Panel B: A one message threshold is observed in three evolutionarily-divergent organisms**. As predicted by the RLTO model, essential, but not nonessential genes, are observed to be expressed above a one message per cell-cycle threshold. All organisms have roughly similar distributions of message number for essential genes, which are not observed for message numbers below a couple per cell cycle. **Panel C: The distribution of gene transcription rate**. The typical transcription rate varies by two orders-of-magnitude between organisms. **Panel D: The distribution of gene cellular message number**. There is also a two-order-of-magnitude variation between typical cellular message numbers. No consistent lower threshold is observed for either statistic.

### 2. Methods: Quantitative estimates of central dogma parameters

#### a. Estimating the cellular message number: μ_m/c_

For each model organism (and condition), we found a consensus estimate from the literature for the total number of mRNA messages per cell 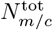. This number and its source are provided in Tab. S2. To estimate the number of messages corresponding to gene *i*, we re-scaled the un-normalized abundance level *r*_*i*_:

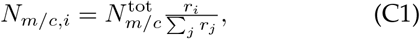

where the sum over gene index *j* runs over all genes.

#### b. Estimating the transcription rate: β_m_

To estimate the transcription rate for gene *i*, we start from the estimated cellular message number *N*_*m/c,i*_ and use the canonical steady-state noise model prediction for the cellular message number:

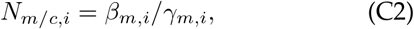

where *γ*_*m,i*_ is the message decay rate. Since gene-to-gene variation in message number is dominated by the transcription rate (*e*.*g* [36]), we estimate the decay rate as the inverse gene-median message lifetime:

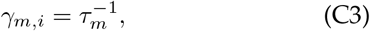

for which a consensus value was found from the literature. This number and its source are provided in Tab. S2. We then estimate the gene-specific transcription rate:

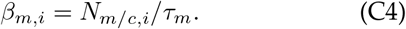

#### c. Estimating the message number: μ_m_

To estimate the message number of gene *i*, we use the predicted value from the canonical steady-state noise model (see A 1 a):

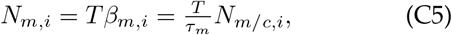

where *T* is the doubling time and *N*_*m/c,i*_ is the cellular message number (Eq. C1).

### 3. Results: Histograms of central dogma transcriptional statistics

We generated histograms for each of the three transcriptional statistics: transcription rate *β*_*m*_, cellular message number *μ*_*m/c*_, and message number *μ*_*m*_. The histograms for transcription rate and cellular message number do not show a consistent lower limit (as predicted) and are shown in Fig. S9; however, the histogram for message number does show a consistent lower bound for the three model organisms and is shown in Fig. S9B.

### 4. Discussion: *E. coli* essential genes below the one-message-rule threshold

Since our own preferred model system is *E. coli*, we focus here. Our essential gene classification was based on the construction of the Keio knockout library [55]. By this classification, 10 essential genes were below threshold. (See Tab. S3.) Our first step was to determine what fraction of these genes were also classified as essential using transposon-based mutagenesis [64, 65]. Of the 10 initial candidates, only one gene, *ymfK*, was consistently classified as an essential gene in all three studies, and we estimate that its message number is just below the threshold (*μ*_*m*_ = 0.4). *ymfK* is located in the lambdoid prophage element e14 and is annotated as a CI-like repressor which regulates lysis-lysogeny decision [66]. In *λ* phase, the CI repressor represses lytic genes to maintain the lysogenic state. A conserved function for *ymfK* is consistent with it being classified as essential, since its regulation would prevent cell lysis. However, since *ymfK* is a prophage gene, not a host gene, it is not clear that its expression should optimize host fitness, potentially at the expense of phage fitness. In summary, closer inspection of below-threshold essential genes supports the threshold hypothesis.

## Appendix D: Analysis of gene-expression noise

This section provides a detailed development of gene expression noise. We continue the discussion of the model from Sec. A1 that provided a self-contained development of the noise models developed by others which are the input to the RLTO model. Secs. D1-D6 describe the RLTO prediction of non-canonical noise scaling and the test of this model.

### 1. Results: RLTO model predicts non-canonical noise scaling

The predicted scaling of the optimal translation efficiency with message number has many important implications, including on the global characteristics of noise. Based both on theoretical and experimental evidence, it is widely claimed that gene-expression noise should be inversely proportional to protein abundance [9, 67]:

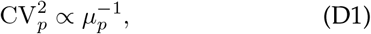

for low-expression proteins, as observed in *E. coli* [9]; however, the more fundamental prediction is that the noise is inversely proportional to the message number:

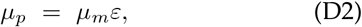

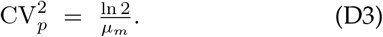

In *E. coli*, the translation efficiency is roughly constant (*i*.*e*.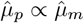, Fig. 4D) and therefore Eq. D3 is consistent with the canonical noise model (Eq. D1). However, in eukaryotes, the translation efficiency grows with message number (*i*.*e*.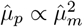, Fig. 4BC). If we substitute this proportionality into Eq. D3, we predict the non-canonical noise scaling:

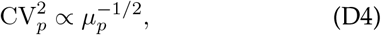

for eukaryotic cells.

### 2. Methods: Analysis of gene expression noise

The quantitative model for gene expression noise includes multiple contributions:

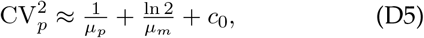

where the first term can be understood to represent the Poisson noise from translation, the second term the Poisson noise from transcription, and the last term, *c*_0_, is called the *noise floor* and is believed to be caused by the cell-to-cell variation in metabolites, ribosomes, and polymerases *etc*. [68, 69].

In the main text of the paper, we have ignored the role of the noise floor in the analysis of noise in yeast. Unlike *E. coli*, where the noise floor is high 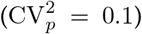 and is determinative of the noise associated with almost all essential genes [9, 68, 69], in yeast the noise floor is much lower 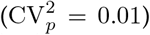 and therefore affects only genes with the highest expression.

**TABLE S3.**
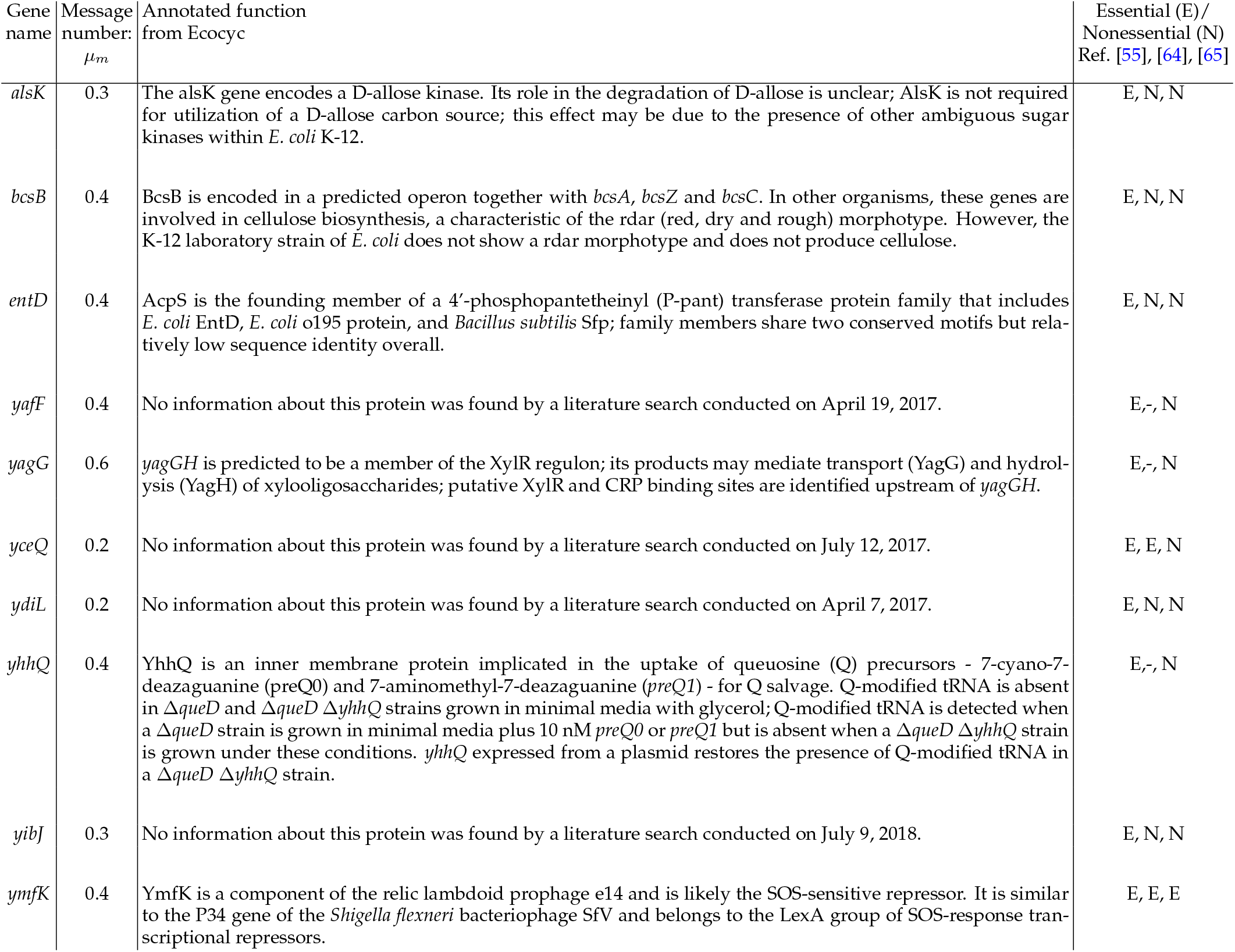
Below-threshold essential genes identified in *E. coli*. This table describes the message numbers and annotations for essential genes that we estimated to have expression below the threshold of one message per cell cycle. However, in the final column, we show classifications from three different studies. Only one of the identified genes, *ymfK*, was consistently defined as essential.

In this section, we will consider models that include the noise floor, since its presence can make the noise scaling more difficult to interpret. To determine if the scaling of the noise is consistent with the canonical assumption that the noise is proportional to 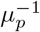 for low expression, we will consider two competing empirical models for the noise (Fig. S10). In the null hypothesis, we will consider a model:

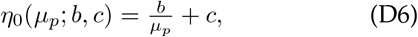

and an alternative hypothesis with an extra exponent parameter *a*:

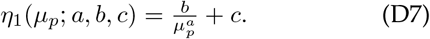

We will assume that 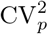 is normally distributed about *η* with unknown variance 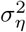.

In this context, a maximum likelihood analysis is equivalent to least-squares analysis. Let the sum of the squares be defined:

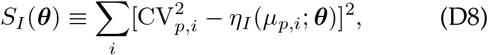

for model *I* where ***θ*** represents the parameter vector. The maximum likelihood parameters are

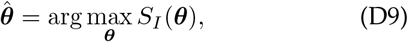

with residual norm:

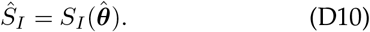

**FIG. S10.**
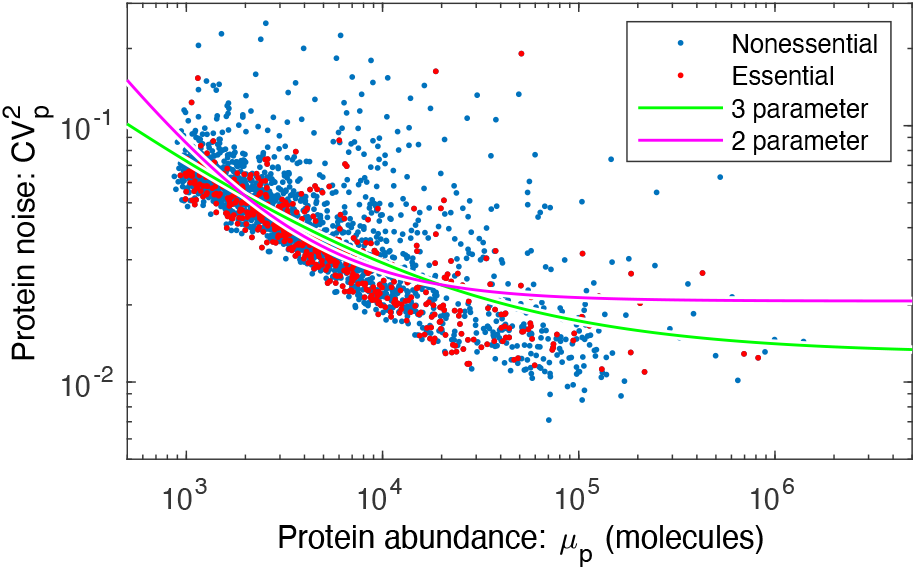
Yeast noise fit against canonical noise model, with a noise floor. Yeast noise data fit with the 2-(null hypothesis with 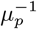 dependence) and 3-parameter 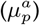 models. The two-parameter model corresponds to the canonical noise model (Eq. D1) and fails to quantitatively fit the data.

To test the null hypothesis, we will use the canonical likelihood ratio test with the test statistic:

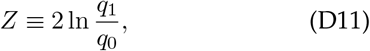

where *q*_0_ and *q*_1_ are the likelihoods of the null and alternative hypotheses, respectively. Wilks’ theorem states that *Z* has a chi-squared distribution of dimension equal to the difference of the dimension of the alternative and null hypotheses (3 *−* 2 = 1).

#### a. Hypothesis test I

In our first analysis, we will estimate the variance directly. We computed the mean-squared difference for successive CV^2^ values, sorted by mean protein number *μ*_*p*_. The variance estimator is

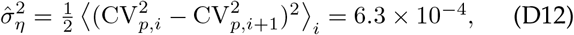

where the brackets represent a standard empirical average over gene *i* for the *μ*_*p*_-ordered gene CV^2^ values. The test statistic can now be expressed in terms of the residual norms:

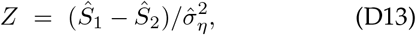

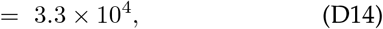

which corresponds to a p-value far below machine precision. We can therefore reject the null hypothesis.

#### b. Hypothesis test II

In a more conservative approach, we can use maximum likelihood estimation to estimate the variance of each model independently as a model parameter. In this case, the test statistic can again be expressed in terms of the residual norms:

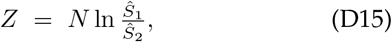

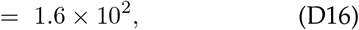

where *N* is the number of data points. (Details of derivation are in Sec. D 2 d.) In this case, the p-value can be computed assuming the Wilks’ theorem (*i*.*e*. the chi-squared test):

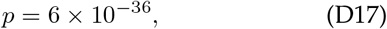

again, strongly rejecting the null hypothesis.

#### c. Maximum likelihood estimates of the parameters

In the alternative hypothesis, the maximum likelihood estimate (MLE) of the empirical noise model (Eq. D7) parameters are (Fig. S10):

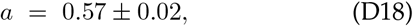

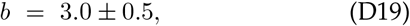

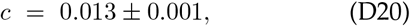

where the parameter uncertainty has been estimated using the Fisher Information in the usual way using the MLE estimate of the variance [70, 71].

#### d. Details: Statistical details MLE estimate of the variance

The minus-log-likelihood for the normal model *I* is:

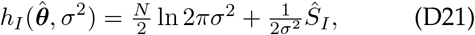

where 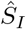 is the least-square residual. We then minimize *h*_*I*_ with respect to the variance *σ*^2^:

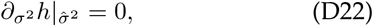

to solve for the MLE 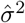:

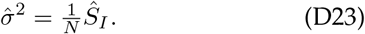

Next we evaluate *h* at the variance estimator:

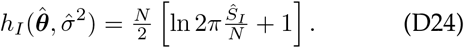

The test statistics can be written in terms of the *h*’s:

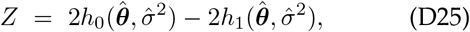

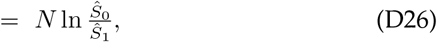

which can be evaluated directly in terms of the residual norms for the null and alternative hypotheses.

### 3. Results: Non-canonical noise scaling is observed in yeast

To test the RLTO model predictions for noise scaling, we reanalyze the dataset collected by Newman *et al*., who performed a single-cell proteomic analysis of yeast by measuring the abundance of fluorescent fusions by flow cytometry [8]. Since the competing models (Eqs. D1 and D4) make different scaling predictions, we first apply a statistical test to determine whether the observed scaling is consistent with the canonical model (Eq. D1). We consider the null hypothesis of the canonical model (Eq. D6) and the alternative hypothesis with an unknown scaling exponent (Eq. D7). To test the models, we perform a null hypothesis test. (A detailed description of the statistical analysis, which includes the contribution of the noise floor, is given in the Sec. D.) We reject the null hypothesis with a p-value of *p* = 6 *×* 10^*−*36^. The observed scaling exponent is *â* = *−*0.57 *±* 0.02, which is close to our predicted estimated exponent from the RLTO model 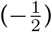.

### 4. Methods: Parameter-free prediction of noise from protein-message relation

By combining the noise model (Eq. D3) with a protein-message abundance relation, the relation between protein abundance and noise can be predicted without additional fitting parameters. (We call this prediction *parameter-free* since, although a parameter is fit when determining the protein-message abundance, once this relation has been established, no new parameters are fit in order to predict the noise.) To test this prediction, we will compare three competing models: (i) the RLTO model, (ii) an empirical protein-message abundance model, and (iii) the constant-translation-efficiency model.

#### a. Estimating protein number (μ_p_) for the noise analysis

The protein abundance data for yeast grown in YEPD media and measured with flow cytometry fluorescence [8] were given in arbitrary units (AU). In order to convert from AU to protein number, the fluorescence values were rescaled by comparing with mass-spectrometry protein abundance data for yeast grown in YNB media [61]. Since the protein abundance from mass-spectrometry was given in terms of intensity, the intensity values were first rescaled by the total number of proteins in yeast, 5 *×* 10^7^. (See Sec. C 1 b.) The massspectrometry protein data was thresholded at 10 proteins, based on the assumption that the noise of the data for 10 and fewer proteins makes the data unreliable. Next, the log of the fluorescence protein abundance in AU as a function of the log of thresholded mass-spectrometry protein abundance was fit as a linear function with an assumed slope of 1 to find the offset, 3.9, (Fig. S11) which corresponds to a multiplicative scaling factor. We then used that offset value to rescale the fluorescence data from AU to protein number. We also compared to yeast grown in SD media [8] and found a similar offset result.

**FIG. S11.**
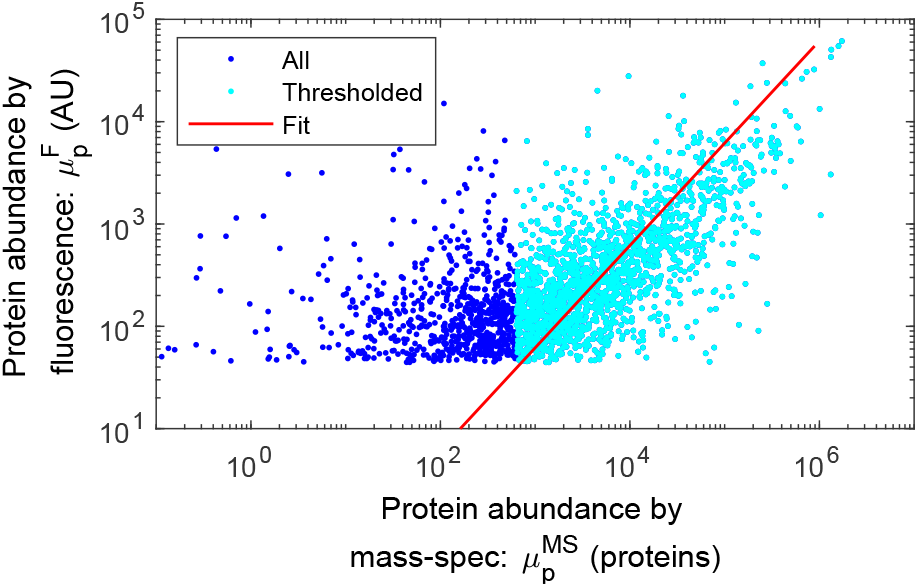
Fit to rescale fluorescence intensity to protein number. Protein abundance from flow cytometry fluorescence [8] as a function of mass-spectrometry scaled abundance [61]. The mass-spectrometry data was thresholded at 10 proteins, and then a linear fit was performed to find the multiplicative offset of 3.9, which was used to convert protein fluorescence AU to number.

#### b. Empirical models for yeast gene expression

To generate the empirical model for protein number as a function of message number, we used protein abundance data from Newman *et al*. [8], re-scaled to estimate protein number (Sec. D 4 a) and transcriptome data from Lahtvee *et al*. [72], re-scaled to estimate message number (Sec. C 2 c).

#### c. Empirical model for protein number

We initially fit the empirical model for protein number,

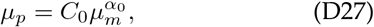

to the data using a standard least-squares approach; however, the algorithm led to a very poor fit since it does not account for uncertainty in both independent and dependent variables. We therefore used an alternative approach [73], which assumes comparable error in both variables. The model parameters are:

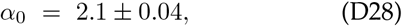

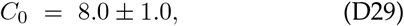

where the uncertainties are the estimated standard errors. The result of the empirical model fit is shown in Fig. S12A, along with the constant-translation-efficiency model, and the RLTO model.

#### d. Empirical model for message number

For the prediction of the coefficient of variation, it is useful to invert Eq. D27 to generate a model for message number as a function of protein number:

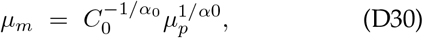

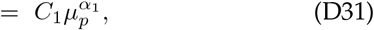

where the last line defines two new parameters: a coefficient *C*_1_ and an exponent *α*_1_. The resulting parameters and uncertainties are:

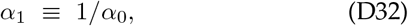

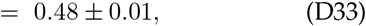

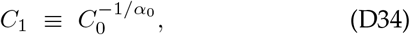

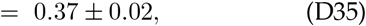

where the uncertainties are the estimated standard errors.

#### e. Empirical model for translation efficiency

To generate an empirical model for translation efficiency, we started from the empirical model for protein number (Eq. D27), and then use Eq. D2 to relate protein number, message number, and translation efficiency:

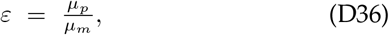

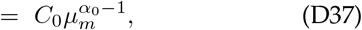

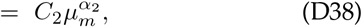

where the last line defines two new parameters: a coefficient *C*_2_ and an exponent *α*_2_. The resulting parameters and uncertainties are:

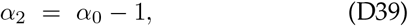

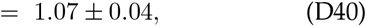

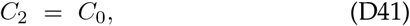

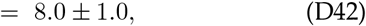

where the uncertainties are the estimated standard errors.

#### f. Empirical model for the coefficient of variation

To generate an empirical model for the coefficient of variation, we started from the empirical model for message number (Eq. D31), and then substitute this into the statistical model prediction for 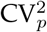 (Eq. D3):

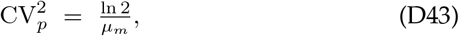

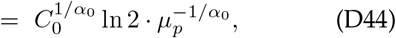

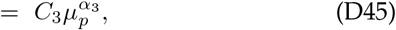

where the last line defines two new parameters: a coefficient *C*_3_ and an exponent *α*_3_. The resulting parameters and uncertainties are:

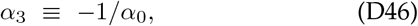

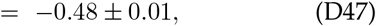

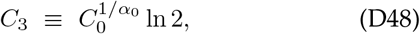

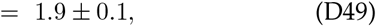

where the uncertainties are the estimated standard errors.

**FIG. S12.**
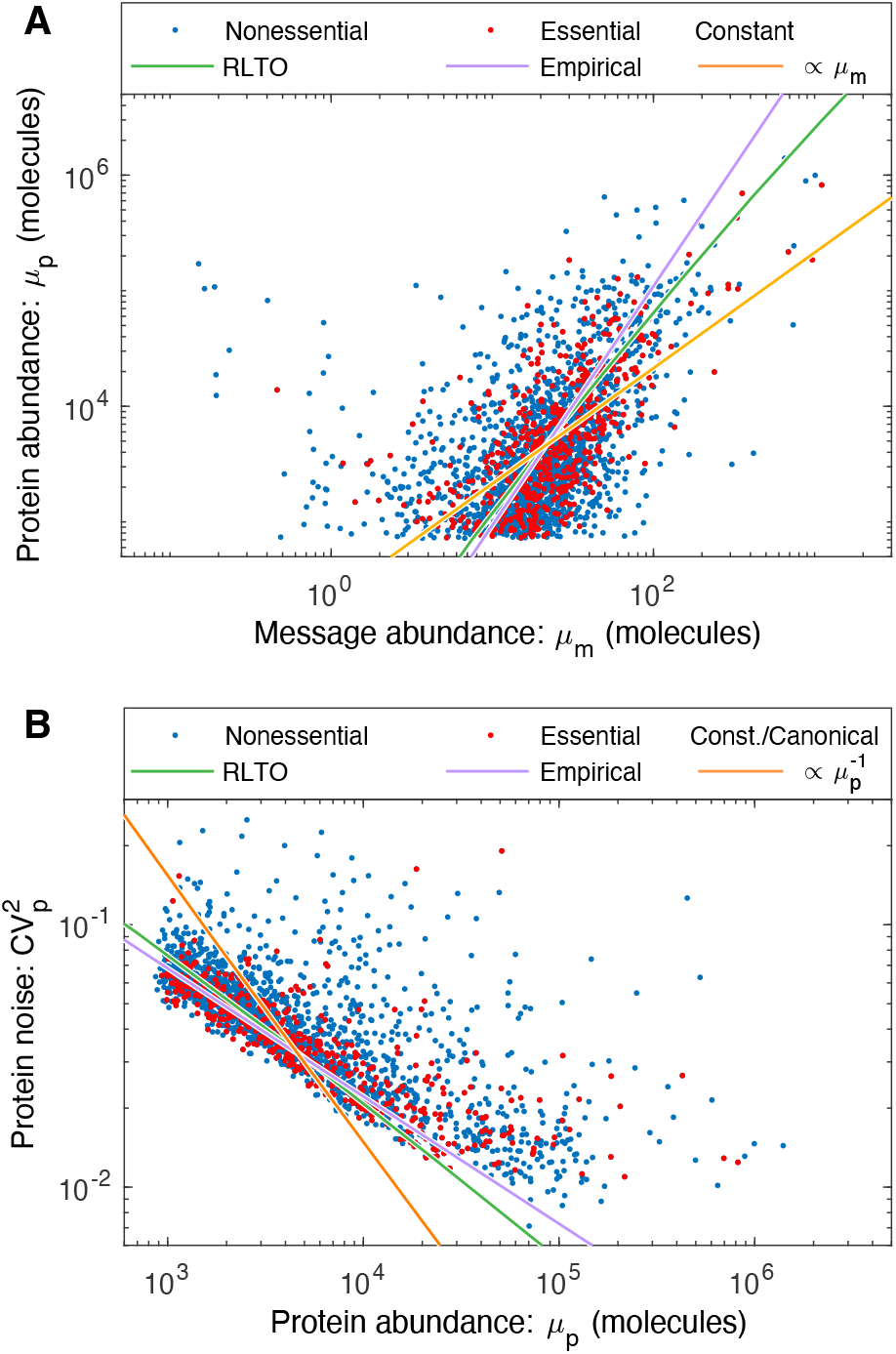
Load balancing predicts the scaling of noise. **Panel A: Three competing models for protein abundance in yeast.** The empirical model (purple) fits the slope and the y offset. The RLTO (green) and constant-translation-efficiency (orange) models fit a parameter corresponding to the y offset only. As discussed in the analysis of the proteome fraction, the RLTO model qualitatively captures the scaling of the protein abundance with message number better than the constant translation efficiency model; however, the predicted fit does not correspond to the optimal power law, which is represented by the empirical model. The protein abundance has a cutoff near 10^1^ due to autofluorescence [8]. **Panel B: Predictions of the noise-protein abundance relation**. Using each competing protein abundance model, the noise-protein abundance relation can be predicted using Eq. D3. The canonical noise model (Eq. D1) fails to capture even the scaling of the noise. In contrast, both the RLTO and empirical models quantitatively predict both the scaling and magnitude of the noise. The empirical model has the highest performance, presumably due to its two-parameter fit to the protein abundance in Panel A. A fit accounting for the noise floor is shown in Fig. S10.

### 15. Results: Parameter-free prediction of noise-abundance in yeast

The fit of the competing protein-message abundance models are shown in Fig. 5A. Using each model, we can now predict the relation between protein abundance and noise without additional fitting parameters. The predictions of the three competing models are compared to the experimental data in Fig. 5B.

In both its ability to capture the protein abundance and predict the noise, the RLTO model vastly outperforms the constant-translation-efficiency model. The purely empirical model that best captures the protein abundance data, due to directly fitting both the y-offset and slope, also performs best in predicting the noise. It is important to emphasize that the prediction of the noise in all models is non-trivial since there are no free parameters fit, once the protein abundance relation is determined. We therefore conclude that the noise model (Eq. D3) quantitatively predicts the observed noise from the message number and that eukaryotic noise has non-canonical scaling due to load balancing.

### 6. Discussion: Implications of noise

What are the biological implications of gene expression noise? Many important proposals have been made, including bet-hedging strategies, the necessity of feed-back in gene regulatory networks, *etc*. [7]. Our model suggests that robustness to noise fundamentally shapes the central dogma regulatory program. With respect to message number, the one-message-rule sets a lower bound on the transcription rate of essential genes. (See Fig. 6B.) With respect to protein expression, robustness to noise has two important implications: Protein over-abundance significantly increases protein levels above what would be required in the absence of noise and therefore reshapes the metabolic budget. (See Fig. 6A.) Robustness to noise also gives rise to load balancing, the proportionality of the optimal transcription and translation rates. (See Fig. 6C.) Not only does robustness to noise affect central dogma regulation, but there is an important reciprocal effect: Load balancing changes the global scaling relation between noise and protein abundance. (See Fig. S12B.)

## Appendix E: Data Tables

### 1. Datasets for one-message-rule analysis

**Data S1**: Organism: *S. cerevisiae* (yeast). Original source: [72], [56], [61] & [8]. Essential gene classification: [19]. Processing: We merged these datasets. We added message number (messages per cell-cycle) as described in Sec. C.

**Data S2**: Organism: *Homo sapiens* (human). Original sources: mRNA abundances: Human Protein Atlas [62]. Essential gene classification: [53]. Processing: We merged these datasets. We added message number (messages per cell-cycle) as described in Sec. C.

**Data S3**: Organism: *E. coli* grown in rich media (LB). Original sources: mRNA abundance: [46]. Essential gene classification: [55]. Processing: We merged these datasets. We added message number (messages per cell-cycle) as described in Sec. C.

**Data S4**: Organism: *E. coli* grown in minimal media (MM), Original sources: mRNA abundance: [46]. Essential gene classification: [55]. Processing: We merged these datasets. We added message number (messages per cell-cycle) as described in Sec. C.

### 2. Datasets for load-balancing analysis

**Data S5**: Load-balancing data for yeast cells. Organism: *S. cerevisiae* (yeast). Original sources: Protein abundance [21]. mRNA abundance: [72]. Processing: We merged these datasets. We added protein number and message number (messages per cell-cycle) as described in Sec. C.

**Data S6**: Load-balancing data for mammalian cells. Organism: *Mus musculus* (mammalian). Original source: [23]. Processing: We added protein number and message number (messages per cell-cycle) as described in Sec. C.

**Data S7**: Load-balancing data for *E. coli*. Organism: *E. coli*. Original source: [22]. Processing: We added protein number and message number (messages per cell-cycle) as described in Sec. C.

### 3. Datasets for noise analysis

**Data S8**: Noise data for yeast. Organism: *S. cerevisiae* (yeast). Original sources: Noise measurements: [8]. Essential gene classifications: [19]. Protein fluorescence data, rescaled by mass-spec data as described in Sec. D 4 a: [8]. Processing: We merged these datasets. We added protein number as described in Sec. C.

### 4. Datasets for one-message-rule exceptions

**Data S9**: Exceptions to the one-message-rule for yeast cells. Organism: *S. cerevisiae* (yeast). Original sources: Message abundances: [72]. Essential gene classifications: [19].

**Data S10**: Exceptions to the one-message-rule for human cells. Organism: *H. sapiens* (human). Original sources: Message abundances: [62]. Essential gene classifications: [53].

**Data S11**: Exceptions to the one-message-rule for *E. coli*grown in rich media (LB). Organism: *E. coli*. Original sources: Message abundances: [46]. Essential gene classifications: [55].

## Notes

### Competing Interest Statement

The authors have declared no competing interest.

### Summary of Updates

Slight phrasing changes; add a few paragraphs to Discussion, fixed some typos; add Supplemental data table descriptions

